# TBX5 dosage governs ventricular cardiomyocyte maturation, specialization and dedifferentiation *in vivo*

**DOI:** 10.64898/2026.04.13.718086

**Authors:** Alexandra E. Giovou, Otto J. Mulleners, Rocco Caliandro, Vincent A.J. Warnaar, Arie R. Boender, Mischa Klerk, Mathilde R. Rivaud, Diederik W.D. Kuster, Mauro Giacca, Bjarke Jensen, Gerard J.J. Boink, Monika M. Gladka, Vincent M. Christoffels

## Abstract

Variation in transcription factor (TF) activity modulates traits and disease susceptibility, yet how such variation translates into cellular phenotype and organ function *in vivo* is not well established. We utilized AAV-mediated gene delivery to express the dosage-sensitive TF TBX5 across a physiologically plausible range in postnatal ventricular cardiomyocytes. Transcriptomic profiling revealed that TBX5 dosage-dependent cardiomyocyte states changed gradually and often non-monotonically across the TBX5 dosage spectrum. The lowest dosages induced cardiomyocyte hypertrophy and upregulated gene programs governing oxidative metabolism, calcium handling, and contractility. In contrast, mid-to-high dosages induced a ventricular conduction system-like transcriptional profile. Supraphysiological dosages triggered cardiomyocyte dedifferentiation, characterized by cardiomyocyte size reduction, cell cycle re-entry, metabolic reprogramming, and impaired ventricular ejection fraction. Furthermore, the ventricular state of an *Nppa-Nppb* deficiency model characterized by cardiac hypertrophy and reduced *Tbx5* expression was partially normalized by TBX5 delivery. By defining the non-linear relationship of TBX5 activity level, cardiomyocyte state and cardiac function, our study reveals how TF dosage regulates the transitions from physiological maturation to specialized lineage acquisition and dedifferentiation at the cell and organ level *in vivo*.

## Introduction

Cellular identity and state are governed by sequence-specific, dosage-sensitive transcription factors (TFs), pivotal components of gene regulatory programs that modulate target gene expression ^1,2^. The impact of TF dosage on molecular and cellular states is well-documented ^3–11^, with TF haploinsufficiency notably enriched among genes linked to developmental defects and disease ^12,13^. Genome-wide association studies (GWAS) have identified numerous trait-associated variants likely affecting TF binding sites ^14–17^, while population genetic studies have revealed that variation in TF loci and tissue-specific expression levels correlates with disease risk ^10,18–22^.

Understanding how phenotypes and organ function respond to shifting TF levels is fundamental. Most functional studies, however, rely on knockout, knockdown, or extreme overexpression models, which do not capture the continuous dosage-to-phenotype relationships underlying complex traits and disease ^3,10^. Recent *in vitro* cell-model studies enabling precise TF manipulation and transcriptomic profiling have begun to identify the intrinsic properties, such as cofactor availability and regulatory DNA composition, that define these responses ^3,5,8,10,23^. Nevertheless, insight into these relationships *in vivo* remains limited, particularly regarding variations across physiologically relevant levels.

Cardiac development and homeostasis depend on the evolutionarily conserved T-box factor TBX5 ^24^. Pathogenic loss-of-function variants in a single TBX5 allele cause Holt-Oram syndrome, characterized by congenital heart defects and conduction abnormalities ^25–28^. Mouse and human stem cell models of TBX5 haploinsufficiency have elucidated its role in morphogenesis, postnatal identity, and ventricular function ^7,9,29–36^. Emerging evidence suggests that increased dosage is equally critical. TBX5 duplications and gain-of-function variants (e.g., G125R) also result in Holt-Oram-syndrome-like phenotypes ^37–40^. Furthermore, GWAS link modestly elevated cardiac TBX5 expression to increased atrial fibrillation risk ^19,21^, a finding supported by mouse models where a 25% increase in *Tbx5* expression induces arrhythmia vulnerability ^41^. These observations underscore that cardiac health requires an optimal TBX5 dosage window.

Here, we investigated the response of the heart to gradual increases in TBX5 dosage. Focusing on postnatal ventricular cardiomyocytes (CMs), non-mitotic cells that maintain an optimal state for lifelong function, we hypothesized that their stability is sensitive to precisely titrated TBX5 levels. Using CM-specific AAV9 vectors to express TBX5 across a physiologically plausible gradient *in vivo*, we found that the transcriptional state and ventricular function are highly dosage-sensitive. Our analysis revealed that these responses are often non-linear and non-monotonic, affecting critical programs such as energy metabolism, conductivity, and calcium handling. Finally, we demonstrate that TBX5 transduction can partially rescue the transcriptional and hypertrophic state associated with *Nppa-Nppb* deficiency. Our results provide key insights into how continuous TF dosage gradients shape the phenotypes underlying complex traits and disease risk.

## Methods

### Animal experiments

Animal care and experiments were performed according to the guidelines from Directive 2010/63/EU of the European Parliament on the protection of animals used for scientific purposes. Animal experiments were approved by the institutional policies and regulations of the Amsterdam University Medical Centers (#AVD11400202216572) in compliance with the Dutch government guidelines. The *Nppa-Nppb*^⁻/⁻^ mouse line has been previously described and characterized. ^42^.

### AAV vectors

| Serotype | Promoter | 5' UTR | Trans gene | Peptide | Nuclear tag | Reporter gene | Name |
| --- | --- | --- | --- | --- | --- | --- | --- |
| AAV6 | CMV | - | - | - | - | GFP | AAV6-Control |
| AAV6 | hTNNT2 | uORF | FLAG-TBX5 | P2A | H2B | GFP | AAV6-Low-TBX5-PHG |
| AAV6 | hTNNT2 | - | FLAG-TBX5 | P2A | H2B | GFP | AAV6-TBX5-PHG |
| AAV9 | hTNNT2 | - | mCherry | P2A | H2B | GFP | AAV9-Control |
| AAV9 | hTNNT2 | uORF | FLAG-TBX5 | P2A | H2B | GFP | AAV9-Low-TBX5-PHG |
| AAV9 | hTNNT2 | - | FLAG-TBX5 | P2A | H2B | GFP | AAV9-TBX5-PHG |
| AAV9 | hTNNT2 | - | FLAG-TBX5 | - | - | - | AAV9-TBX5 |
| AAV9 | hTNNT2 | - | - | - | - | GFP | AAV9-GFP |

### AAV vector production

The AAV9-TBX5 and AAV9-GFP were produced by the AAV Vector unit at ICGEB Trieste. For all the remaining vectors: AAV6 vectors were produced by double transfection of HEK293T cells and AAV9 vectors by triple transfections. Low passage HEK293T cells were plated in thirty 145 mm dishes at the density of 1.5×10^7^ cells per dish in DMEM-GlutaMax (ThermoFisher Scientific) containing 10% fetal bovine serum (FBS) (Sigma) and 1% penicillin-streptomycin (P/S) (ThermoFisher Scientific). For AAV6 vectors, cells were transfected the next day with 33 µg pDP6 and 16 µg AAV transfer plasmids per dish using linear polyethylenimine (PEI) (Polysciences Inc). For AAV9 vectors, cells were transfected with 13 µg Rep2Cap9 plasmid, 21 µg helper plasmid, and 13 µg AAV transfer plasmid per dish. Medium was also replaced during transfection to DMEM containing 1% P/S. Three days after transfection, cells were collected by centrifugation, and the medium was concentrated by tangential flow filtration using the ÄKTA Flux S system (GE Healthcare) to a final volume of 20 ml. Cells and medium were then combined, frozen and thawed twice followed by DNAseI, Rnase A, and Benzonase treatment. AAV vectors were purified by iodixanol density-gradient ultracentrifugation overnight. The AAV-containing fraction was then collected and concentrated to 1 mL by buffer exchange to PBS containing 0.001% Pluronic F68 using Amicon Ultra-15 100 kDa centrifugal filter units (Millipore). Concentrated AAV vectors were aliquoted and stored at −80°C until use. Genomic titer was determined by qPCR.

### *In vivo* AAV9 transduction

For the baseline experiments, juvenile FVB/NJ mice (postnatal day 15, P15) were injected retro-orbitally with 50ul of PBS containing 1×10^11^ vg/mouse using a 0.3 mm G U-100 insulin syringe (BD micro-Fine). Adult *Nppa-Nppb*^⁻/⁻^ mice (8 weeks old) at baseline were retro-orbitally injected with 50 ul of PBS containing 1×10^12^ vg/mouse. For all experiments both male and female mice were used.

### EdU injections in mice

To assess cell cycle re-entry mice received bi-daily subcutaneous injections of 5-ethynyl-2’-deoxyuridine (EdU) (Thermofisher) diluted in PBS using a 0.3 mm G U-100 insulin syringe (BD micro-Fine). EdU concentration was delivered at 50 mg/kg diluted in PBS in 50 µl and 100 µl volume for the juvenile and adult mice, respectively.

### Heart collection and histological analysis

Hearts were excised, washed in PBS and fixed in 4% PFA at room temperature (on a shaker) for 24 hours before exchanging to 70% ethanol. The paraffin-embedded hearts were sectioned at 5 µm and mounted on microscope slides before deparaffinization and rehydration in an ethanol gradient. Hematoxylin and Eosin and picrosirius red stainings were used to assess cardiac morphology and fibrosis in the juvenile hearts, respectively. The heart and tibia were collected at the end of each experiment to assess heart weight and tibia length ratios.

### Immunohistochemistry

Histological sections were boiled in antigen retrieval solution for 5 min at 1000 W and rinsed in Demi water followed by PBS for 5 min. Sections were treated with 0.1% Triton-X/PBS for 8 min. Blocking was performed with 4% BSA for 30 min. EdU was visualized with a Click-iT EdU Cell Proliferation Imaging Kit, Alexa Fluor 647 (Thermo Fisher Scientific, C10340), according to the manufacturer’s instructions. Sections were incubated overnight at 4°C with primary antibodies including chicken anti-GFP (AvesLab, #GFP-1020, 1:400), mouse anti-TNNT2 (Thermo Fisher Scientific, MA512960 1:200), rabbit anti-DMD (Abcam, Ab15277, 1:200), mouse anti-KI67 (BD Pharma/Becton&Dickinson, 566003, 1:200), goat anti-GJA5 (Santa Cruz/Tebu-bio, sc-20466, 1:150), rabbit anti-NPPA (Campro SC, RGG9103, 1:200), rabbit anti-PCM1 (Atlas antibodies/Bio-connect, HPA023370), goat anti-TBX5 (Santa Cruz/Tebu-bio, sc-17866, 1:200). Afterwards, sections were washed 3 times for 5 min in PBS/tween and incubated at room temperature for 2 hours with corresponding secondary antibodies Alexa Fluor 647 donkey anti-rabbit (Invitrogen, #A31573, 1:250), Alexa Fluor 647 donkey anti-goat (Invitrogen, #A-21447, 1:250), Alexa Fluor 647 donkey anti-mouse (Invitrogen, #A-231571, 1:250), Alexa Fluor 555 donkey anti-rabbit (Invitrogen, #A31572, 1:250), Alexa Fluor 555 donkey anti-mouse (Invitrogen, #A31570, 1:250), Alexa Fluor 488 donkey anti-chicken (Jackson ImmunoResesearch, #134115, 1:250) and DAPI (Serva, #18860, 1:1000). Following secondary antibody incubation, sections were washed 3 times for 5 min in 0.05% Tween-20/PBS and mounted using 50:50 % glycerol/PBS. Sections stained for NPPA were not boiled in antigen retrieval, but the rest of the protocol was followed accordingly. For quantification of GFP, EdU, KI67, and DMD, images were acquired under a Leica DM6000 microscope using 20× objectives. Confocal images were acquired using Leica TCS SP8 X White laser or a Stellaris 5 confocal microscope with a 40× objective.

### Quantification of cell size and cell cycle reactivation

Cardiomyocyte (CM) cell size was quantified manually on three sections per heart (spaced at 1:30 intervals) using DMD⁺/DAPI⁺ nuclei as markers (60 CMs per section), in a blinded manner. For each section, a plane close to the luminar side of the heart was captured to ensure equal CM orientation. EdU and KI67 quantification was performed blindly over 1 section/ heart. All analysis was performed with ImageJ software. For quantification of EdU, KI67, and DMD images were acquired under a Leica DM6000 microscope using 20× objective.

### Quantification of nuclear TBX5 intensity

The relative expression of nuclear TBX5 intensity was quantified on sections from mice injected with AAV9-Control, AAV9-Low-TBX5-PHG and AAV9-TBX5-PHG by using Fiji ^43^. Atrial TBX5 nuclei intensity of the AAV9-Control was quantified to establish the physiological ranges of the TBX5 expression. Segmentation of the nuclei was performed using StarDist 2D (v0.03) on the DAPI channel ^44^. The ROIs were then applied on the TBX5 channel, and the mean gray value was measured for the quantification of the TBX5 nuclear signal. Background correction was performed on ventricular control sections, not expressing TBX5 and then consistently applied across all measurements. Nuclear TBX5 intensity was quantified per condition with the following nuclei counts: Atrial AAV9-Control: 673 nuclei; Ventricle AAV9-Control: 980 nuclei; AAV9-Low-TBX5-PHG: 873 nuclei; AAV9-TBX5-PHG: 1179 nuclei (n=3 hearts per condition). Confocal images were acquired with a Leica Stellaris 5 confocal microscope.

### RNA isolation and qPCR

Total RNA was isolated from mouse hearts using ReliaPrep™ RNA Miniprep Systems (Promega, #Z6012) according to the manufacturer’s protocol.RNA-sequencing and gene ontology

750-1500 ng of total RNA was delivered to the Core Genomics Facility of Amsterdam UMC for library preparation and RNA sequencing. Tapestation assessment was performed to ensure the RNA Integrity Number (RIN) of the RNA. RNA Library Prep was performed using KAPA Hyperprep with RiboErase (Roche) according to the manufacturer’s instructions. Paired-end RNA sequencing was performed on the NovaSeq 6000 S4 platform (Illumina). Approximately 40-50 million reads/samples were retrieved. Data analysis was performed by uploading Fastq files to the Galaxy platform ^45^. Fastq files were trimmed using the algorithm “Trim Galore” and mapped on the GRCm38/mm10 mouse genome. DESeq2 (v1.40.2) analysis was performed across groups to identify significantly up- or downregulated genes and to generate normalized counts used for Z-score assessment across groups ^46^. Gene Ontology (GO) analysis and KEGG Pathways analysis of differentially expressed genes (DEGs) were performed using Database for Annotation, Visualization, and Integrated Discovery ^47^.

### Nuclei isolation

For isolation of nuclei, a hybrid isolation protocol involving a sucrose gradient and FACS sorting clean-up was followed. In brief, approximately 70 mg of mouse tissue was homogenized in ice-cold LYS buffer (10 mM Tris-HCl pH 8.0; 5 mM CaCl2; 2 mM EDTA; 0.5 mM EGTA; 1 mM DTT; 3 mM Mg(CH₃COO)₂, 100 U/mL murine RNAse inhibitors) supplemented with 0.4% v/v Triton X-100 using first a tissue homogenizer (IKA, #0003737000) and a glass douncer, subsequently. The tissue homogenate was then kept under constant shaking (35 rpm) on a rotating tube mixer for 10 minutes at 4°C. One volume of ice-cold LYS buffer was added to the tissue homogenate before passing it through a 100µm and 30µm cell strainer. Cell-strained tissue homogenate was centrifuged at 1000g for 5 min at 4°C, and pellet of nuclei and small tissue debris was resuspended in G30 solution (30% iodixanol, 25 mM KCl, 5 mM MgCl2, 20 mM Tricine-KOH, pH 7.8). G30 containing nuclei and tissue debris was layered on top of a preformed density gradient of G30 and centrifuged at 8000g for 20 min at 4°C. Pellet of nuclei was washed in RES buffer (5% BSA-PBS) and resuspended in RES buffer supplemented with 150 U/mL murine RNAse inhibitors. Following homogenization, nuclei were stained with DAPI (ThermoFisher, #62248). Approximately 150,000 DAPI+ events were sorted (Sony SH800 cell Sorter, Sony Biotechnology) in RES buffer. FACS sorting gates were set using nuclei samples resuspended in RES buffer only (no DAPI). Next, sorted nuclei were centrifuged at 1000g for 5 min at 4°C, and the total volume was reduced by approximately ¾ to ensure proper nuclei/μL concentration. Finally, nuclei concentration was determined using Countess II automated cell counter (Invitrogen, AMQAX1000).

### Single-nuclei RNA (snRNA) sequencing

Nuclei were obtained from four pooled hearts per treatment condition; 2 females and 2 males per condition. A volume of nuclei solution containing approximately 33,000 nuclei/sample was loaded on ChromiumX (10x Genomics) for single-nuclei tagging and library preparation (GEM-X Single Cell 3’ Expression Library v3, 10X Genomics). cDNA quality assessment was performed using D5000 ScreenTape Assay on 4200 TapeStation (Agilent). All samples passed the quality control and were sequenced on NovaSeqXPlus (Illumina) with a depth of sequencing of 80 million paired-end reads per sample.

### Single-nucleus RNA-sequencing processing and analysis

Raw sequencing data were processed using Cell Ranger (v9.0.1) ^48^ to generate nucleus-gene count matrices, which were imported into R (v4.4.2) and analyzed using Seurat (v5.3.0, R package) ^49^. Low-quality nuclei were filtered using the following thresholds: nUMI ≥ 800, nUMI ≤ 7,000, nGene ≥ 600, log₁₀(nGene/nUMI) ≥ 0.9, and mitochondrial ratio (mitoRatio) < 0.02. Gene expression data were normalized and variable features identified with FindVariableFeatures. Principal component analysis (PCA) was performed on the scaled expression matrix, retaining components until cumulative variance exceeded 90% and incremental variance contribution fell below 5%. These principal components were used for nearest-neighbor graph construction (FindNeighbors) and UMAP embedding, followed by Louvain clustering (FindClusters, resolution = 0.4).

Potential doublets were identified and removed using DoubletFinder (v2.0.6, R package) ^50^. Artificial doublets were merged with real nuclei, and the optimal pK parameter was determined by maximizing the bimodality coefficient (BCmvn) distribution, yielding pK = 0.23. The expected doublet count was estimated using a multiplet rate of 0.12, based on manufacturer guidelines (Chromium Next GEM Single Cell 3’ Reagent Kits v3.1, Document Number CG000315 Rev F, 10x Genomics, (2024, April), and adjusted for homotypic doublets using modelHomotypic (DoubletFinder). After doublet removal, expression values were normalized using SCTransform (Seurat v5.3.0, R package) ^51,52^ for variance stabilization, applied separately to each sample without dataset integration. Sex assignment was inferred using the classifySex function from speckle (v0.03, R package) ^53^, which classifies nuclei based on the expression of X- and Y-linked genes. A female bias was observed in the AAV9-Low-TBX5-PHG treatment group. As our bulk RNA-seq experiment with the same design showed no sex-specific effects (data not shown), the top 300 most sex-biased genes (Padj < 0.05, |log₂FC| > 1, ranked by |log₂FC|) identified using FindMarkers (Seurat) were removed to avoid confounding, primarily affecting sex-linked transcripts.

Cell types were first assigned using Azimuth (v0.5.0, R package) ^54^ to obtain coarse reference-based annotations. These assignments were refined using FindAllMarkers (Seurat) and known marker genes of major cardiac cell types, identifying principal populations including CM, endothelial cells, fibroblasts, and immune cells. To facilitate cross-species comparisons, gene identifiers were converted to human orthologs using babelgene (v22.9, R package) ^55^. Each major cell type was subsequently subset and reprocessed using the same SCTransform workflow to increase within-type resolution, particularly for CM subclusters. Differential gene expression analyses were performed using FindMarkers (Seurat; Wilcoxon rank-sum test, false discovery rate (FDR)-Padj < 0.05). Because multiple animals were pooled without individual barcodes, pseudo-bulk aggregation was not possible, and nuclei were treated as independent observations within conditions. Differences in the proportions of cell types and CM subclusters across treatments were evaluated using chi-square tests on contingency tables, with significance accepted at p < 0.05 after multiple-testing correction. To compare transcriptional differences between clusters and visualize dose-dependent responses, nuclei were randomly downsampled to 500 cells per cluster. Expression values were converted to z-scores per gene, using genes previously identified from bulk RNA-seq as dosage-responsive (**Figure 2k**; **Figure 6c-f**). Hierarchical clustering was performed in R using dist and hclust (base R), applied to the rows (genes), while nuclei were clustered within their main cluster to preserve cell-type contiguity. The resulting z-score matrix was visualized using ComplexHeatmap (v2.24.1, R package) ^56^, showing hierarchically clustered genes on the rows and ordered nuclei on the columns. Pseudotime trajectories were reconstructed using Slingshot (v2.16, R package) ^57^. Analysis was restricted to CM clusters CM1 to CM6, excluding uCM and VCS. Pseudotime was inferred from the UMAP embedding using the identified Louvain clusters, with CM1 defined as the starting cluster and CM6 as the terminal cluster. To identify genes with dynamic expression patterns along pseudotime trajectories, we applied Generalized Additive Model (GAM)-based testing using tradeSeq (v1.22.0, R package), fitting smooth functions with natural cubic splines (k = 6 knots) and assessing significance with the associationTest function (FDR < 0.05) ^58^. Expression values were averaged across 50 fixed pseudotime bins and visualized using ComplexHeatmap, with hierarchical clustering (Euclidean distance, Ward’s method) applied to the genes. The dendrogram was cut into six clusters, and clusterProfiler (v4.16.0, R package) ^59^, was used to perform Gene Ontology Biological Process enrichment analysis (Padj < 0.05), identifying pathways associated with progressive transcriptional transitions along the CM trajectory.

### Functional annotation

GO enrichment results were filtered prior to term selection to remove poorly informative categories. GO terms were filtered based on the number of genes annotated to the term; only terms with greater than 1 and fewer than 1500 annotated genes were retained. In addition, only GO terms with greater than 1 gene from the queried set assigned to the term were kept. Finally, only terms with a nominal p-value below 0.05 were retained. Redundant terms were subsequently collapsed using the simplify function from the clusterProfiler package with a semantic similarity cutoff of 0.7.

For **Supplementary Figure 2b-c**, 10 GO terms per direction (up- and downregulated) were selected exclusively by keyword-based filtering of GO term descriptions to identify cardiac-relevant functional processes, including contractile function, electrophysiology, metabolism, cellular organization, and developmental or regulatory processes. The keyword list used for filtering was: *heart, cardiac, muscle, contraction, sarcomere, action potential, ion transport, calcium, membrane potential, metabolic, mitochondrial, oxidative, ATP, respiratory, fatty, lipid, cytoskeleton, actin, myofibril, and filament*. All matching terms were ranked by adjusted p-value and the top 10 terms per direction were selected.

For **Figure 5h**, 15 GO terms per direction were selected using a two-step approach. First, the 10 terms with the lowest adjusted p-values were chosen. Second, up to 5 additional terms were selected by keyword-based filtering using the same keyword list as above, ranked by adjusted p-value, excluding any terms already present in the top 10, until a total of 15 terms per direction was reached.

### Mitochondrial to genomic DNA ratio

Quantification of mtDNA copy number was performed as previously described ^60^. Briefly, equal amounts of frozen ventricular tissue (10 mg) from either uninjected or AAV9-TBX5 injected mice was added to 300 µl lysis buffer containing 3 µl proteinase K. Samples were incubated at 55°C overnight. Afterwards, the samples were treated with RNase A (100 µg/ml) and incubated at 37°C for 30 min. Next, the samples were treated with 250 µl ammonium acetate (7.5M) and 600µl of isopropanol, mixed thoroughly and centrifuged at 15,000 g at 4°C for 10 min. After removing the supernatant, the pellet was washed with 500 µl of 70% ethanol. After airdrying, the pellet was resuspended in 100µl TE buffer. Samples were further used for qPCR. Quantitative PCR (qPCR) was performed on LightCycler 480 Instrument II (Roche, #05015243001) using LightCycler 480 SYBR Green I Master (Roche, #04707516001). Primer sequences are shown in **Supplementary File, Table S1**.

### Mouse echocardiography

Cardiac function was evaluated by echocardiography 2 weeks after delivery of the AAV9-virus for the baseline experiments. All measurements were performed using a Visual Sonic Ultrasound system with a 30 MHz transducer (VisualSonics Inc.). Mice were sedated (2% isoflurane) and cardiac imaging was performed at the level of the papillary muscles in a parasternal long-axis view to record M-mode measurements, including left ventricular internal diameter and left ventricular posterior wall thickness, and B-mode measurements, including left ventricular end-diastolic volume and left ventricular end-systolic volume. Fractional shortening was calculated as the end-diastolic dimension minus the end-systolic dimensions normalized to the end-diastolic dimension, and ejection fraction was calculated as stroke volume normalized to end-diastolic volume.

### Mouse electrocardiograms

Animals were anesthetized by 4% isoflurane inhalation and maintained under anesthesia with 2% isoflurane in 1 L/min O_2_. Subcutaneous recording electrodes were placed at the left armpit, right armpit and left groin and ECGs were recorded for a period of 1 min. ECG parameters (RR, PR, and QRS intervals, J and T wave amplitudes) were calculated from lead I and lead II using LabChart Pro 8 (ADInstruments).

### Oroboros analysis of oxidative phosphorylation and fatty acid oxidation

#### Tissue preparation

Left ventricular (LV) tissue was cut into smaller pieces and permeabilized in 2 mL of ice-cold BIOPS solution supplemented with 20 µL of saponin (5 mg/mL) for 30 minutes on ice. Following permeabilization, samples were washed twice for 10 minutes in mitochondrial respiration medium (MIR05), containing: sucrose (110 mM), MgCl₂ (3 mM), KH₂PO₄ (10 mM), HEPES (20 mM), EGTA (0.5 mM), taurine (20 mM), and potassium lactobionate (60 mM), with 1 g/L fatty acid-free bovine serum albumin and the pH adjusted to 7.1 using KOH. After washing, tissues were dried, weighed, and transferred into a high-resolution respirometer (Oxygraph-2k; Oroboros Instruments, Innsbruck, Austria). All experiments were conducted at 37°C, maintaining oxygen concentrations above 300 µM throughout to prevent oxygen diffusion limitations. For each sample, both the oxidative phosphorylation and the fatty acid oxidation were measured in duplicate, and results were expressed as the mean.

#### Oxidative phosphorylation measurements

Leak respiration was measured by adding sodium glutamate (10 mM), sodium malate (2 mM), and sodium pyruvate (5 mM) to support electron flow through complex I. NADH-linked respiration was then determined following the addition of ADP (5 mM). To evaluate the integrity of the outer mitochondrial membrane, cytochrome c (10 µM) was added. Subsequently, OXPHOS capacity, supported by both complex I and complex II, was assessed by adding succinate (10 mM). The excess capacity of the electron transport system (complexes I-IV) was determined by stepwise titration with FCCP (0.05 µM). Succinate-linked respiration was then evaluated after inhibition of complex I with rotenone (0.5 µM). Finally, residual oxygen consumption was determined by fully inhibiting mitochondrial respiration with antimycin A (2.5 µM) and subtracted from all values to obtain net respiration rates.

#### Fatty acid oxidation

To assess fatty acid oxidation, mitochondrial octanoylcarnitine oxidation, proceeding independently of the mitochondrial carnitine shuttle, was measured. Prior to substrate addition, a low concentration of malate (0.1 mM) and ADP (5 mM) were added. Octanoylcarnitine (0.2 mM) and cytochrome c (10 µM) were then added, and the resulting increase in oxygen consumption was taken as an indicator of fatty acid oxidation capacity.

### Statistical analysis

Biological replicates or experimental replicates (n) are indicated on each graph and the corresponding figure legend. Data are presented as the mean ± standard deviation (SD), unless stated differently in the figure legend. Mann-Whitney U test was used to statistically compare two groups. One-way ANOVA or Kruskal-Wallis was used to statistically compare three or more groups. Kolmogorov-Smirnov test was used to statistically compare cumulative distribution between conditions. Differences were considered statistically significant at p < 0.05. Statistical analyses were performed using Prism (GraphPad Prism, version 10).

### Data availability

All processed data are available in a supplementary data Excel file, and unprocessed data have been uploaded to GEO with accession numbers GSE320117 and GSE320357. Human bulk RNA-seq data were obtained from the GTEx Portal (https://gtexportal.org/home/downloads/adult-gtex/bulk_tissue_expression), dbGaP accession number phs000424.v10.p2 accessed on 11/12/2025. Previously published *Nppa-Nppb*^⁻/⁻^ RNA-seq data were retrieved from GSE320356 ^61^. Previously published RNA-seq data comparing atrial and ventricular cardiomyocytes were retrieved from GSE215021 ^62^.

## Results

### Establishing a physiologically plausible TBX5 dosage range in ventricular cardiomyocytes *in vivo*

*Tbx5* is expressed predominantly in the cardiomyocytes (CMs) of the developing and mature heart ^24^. To assess the cardiac *Tbx5* mRNA expression profile across different cardiac regions and developmental stages, we used published spatial transcriptomic datasets (**Figure 1a**) ^63^. *Tbx5* expression is highest in adult atrial myocardium. Fetal ventricular trabecular myocardium expresses approximately 3-4 times less than adult atrial levels, while both fetal compact myocardium and adult ventricular myocardium express approximately 8-fold less *Tbx5* than adult atrial tissue. Normalized expression levels of *TBX5* in adult human right atrial and left ventricular tissues (GTEx ^64^) showed relative differences comparable to those in their mouse counterparts (**Figure 1a**). The well-established TBX5 target genes *Nppa* and *Gja5* ^26^ display expression levels that respond proportionally to *Tbx5* dosage. Published scRNA-seq data revealed that within early postnatal ventricles, Purkinje fiber cardiomyocytes of the ventricular conduction system (VCS) express substantially higher levels of *Tbx5* than adjacent ventricular contractile cardiomyocytes (mean normalized expression 0.225 vs 0.073; ∼3-fold enrichment) ^65^. Thus, ventricular *Tbx5* expression in the ventricular CMs is reduced after birth, but is maintained at prenatal levels in the VCS. Of note, in the adult, both the ventricular CMs and the VCS require *Tbx5* expression to maintain heart function ^29,34^.

**Figure 1.**
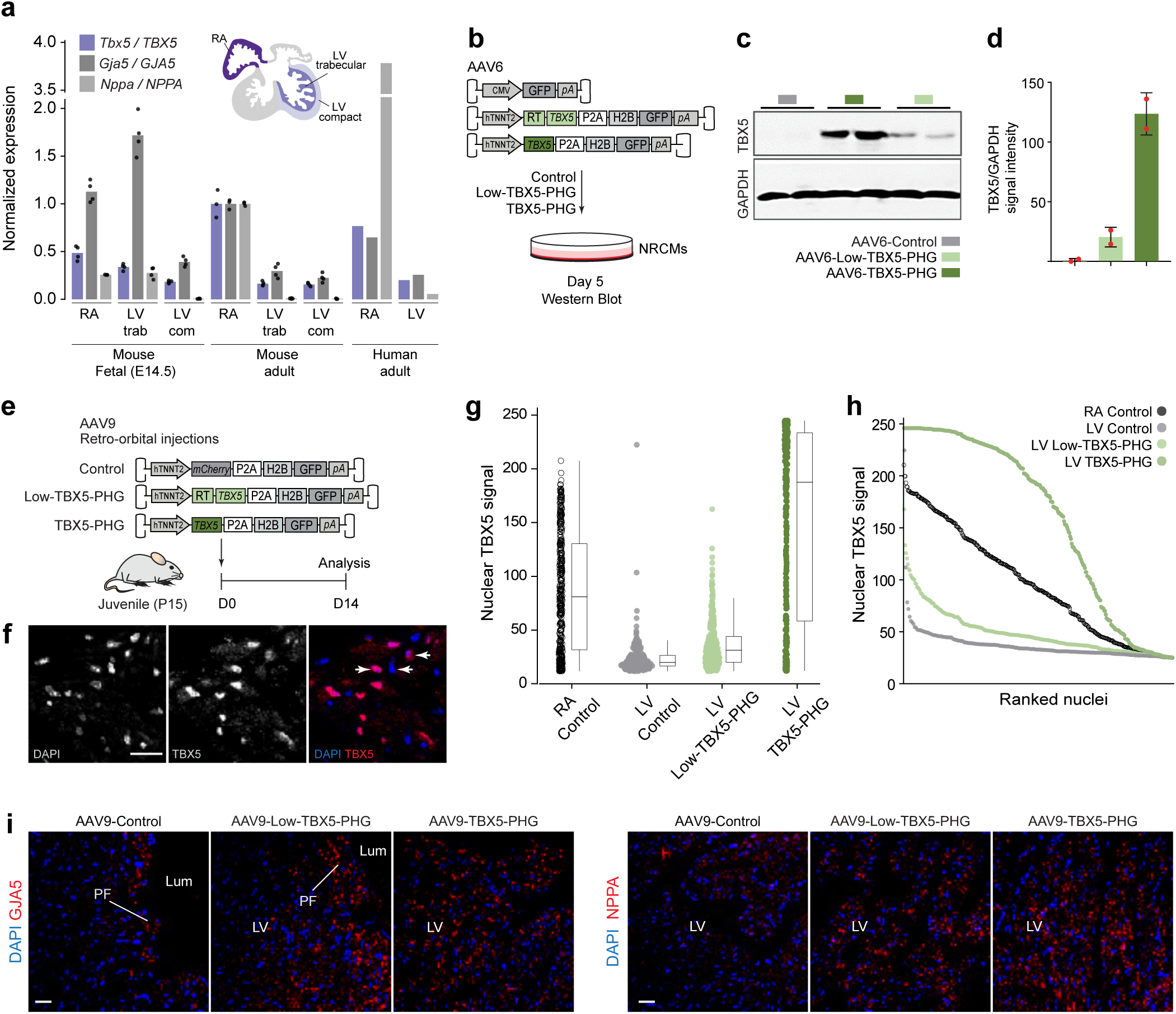
AAV-mediated TBX5 overexpression leads to physiologically relevant TBX5 levels. (**a**) Normalized *Tbx5*/*TBX5, Gja5/GJA5, Nppa/NPPA* expression levels in the right atrium, trabecular left ventricle and compact left ventricle for E14.5 mouse, adult mouse and adult human hearts. Fetal and adult mouse data were obtained from NanoString GeoMx ^63^, human data was retrieved from GTEx, expression levels were normalized to the 300 most stable genes between the three datasets. (**b**) Schematic of *in vitro* experimental model in NRCM, (**c**) Western blot showing TBX5 expression in AAV6-Control, AAV6-Low-TBX5-PHG and AAV6-TBX5-PHG-treated NRCM (n=2), (**d**) Quantification of TBX5 over GAPDH expression (n=2), (**e**) Schematic of *in vivo* experimental model in P15 mice, (**f**) Representative immunofluorescent staining of DAPI (left) and TBX5 (middle; antibody cross-reactive with human and mouse), and the merged image (right; DAPI in blue and TBX5 in red) in AAV9-Control-treated atrium (Scale bar: 20 μm). (**g**) Relative nuclear intensity (arbitrary units) of TBX5 expression levels were quantified on sections of atria and ventricles for the different treatment conditions. (n=3 hearts per condition). (**h**) Ranked distribution of the same nuclear TBX5 intensity data shown in (**g**). Each dot represents an individual nucleus from atrial and ventricular sections, ranked from lowest to highest intensity within each treatment condition. Colors indicate different treatment conditions. (n=3 hearts per condition), (**i**) Representative immunofluorescence staining of DAPI (blue) and GJA5 (red, left row) or NPPA (red, right row) in AAV9-Control-treated ventricle (left), AAV9-Low-TBX5-PHG-treated ventricle (middle), and AAV9-TBX5-PHG-treated ventricle (right) (Scale bar: 20 μm).

We next sought to establish a physiologically relevant TBX5 dosage range in fully differentiated cardiomyocytes. AAV vectors expressing different levels of TBX5 tagged with P2A-H2B-GFP (PHG) allowing visualization of the nuclei of transduced cells were constructed. To reduce TBX5 expression levels from the expression vector by about 5 fold, a small open reading frame (uORF) was inserted in the 5’UTR of the TBX5 coding sequence ^66,67^. To validate reduced protein expression by uORF inclusion, neonatal rat ventricular CMs (NRCM) were transduced with AAV6-CMV-GFP, AAV6-hTNNT2-Low-TBX5-PHG and AAV6-hTNNT2-TBX5-PHG (hereafter referred to as AAV6-Control, AAV6-Low-TBX5-PHG and AAV6-TBX5-PHG, respectively) for 5 days followed by Western blot analysis (**Figure 1b-d**). Next, AAV9-TNNT2-mCherry-PHG (AAV9-Control), AAV9-hTNNT2-Low-TBX5-PHG (AAV9-Low-TBX5-PHG) and AAV9-hTNNT2-TBX5-PHG (AAV9-TBX5-PHG) were systemically injected in postnatal day 15 (P15) mice, and after two weeks, the hearts were collected for analysis (**Figure 1e**).

To determine AAV9-delivered TBX5 expression levels in ventricular myocardium, we measured nuclear TBX5 immunofluorescence signal intensity in confocal sections from AAV9-Control, AAV9-Low-TBX5-PHG and AAV9-TBX5-PHG transduced ventricles. As a reference for maximal endogenous TBX5 expression, we measured nuclear TBX5 signal in postnatal atrial tissue in the Control treated animals, respectively (**Figure 1f**). Across hundreds of nuclei, TBX5 signal intensities covered a broad range from undetectable levels (presumably TBX5-negative nuclei) to intensities exceeding those observed in atrial nuclei (**Figure 1g-h**). Combining AAV9-Control, AAV9-Low-TBX5-PHG, and AAV9-TBX5-PHG conditions, the majority of ventricular nuclei expressed TBX5 at a level range spanning from low endogenous postnatal ventricular levels to high endogenous postnatal atrial levels, a range that encompasses the levels observed in fetal trabecular myocardium. However, in a subset of nuclei, particularly in the AAV9-TBX5-PHG hearts, AAV9-driven TBX5 expression exceeded maximal atrial levels (**Figure 1g-h**). Delivery of AAV9-Low-TBX5-PHG and AAV9-TBX5-PHG resulted in gradual upregulation of TBX5 target gene products GJA5 and NPPA, in the ventricular CMs (**Figure 1i**). Thus, our approach provides a continuous TBX5 dosage range across ventricular CMs *in vivo*, with a portion encompassing physiologically relevant dosages.

### Mapping of the transcriptional states of ventricular CMs across a continuous TBX5 dosage range

We next aimed to investigate the relationship between gradual changes in TBX5 dosage and transcriptional states of CMs at a higher resolution. To this end, we performed single-nucleus RNA sequencing (snRNA-seq) of ventricular tissue (nuclei from four pooled hearts per treatment condition; 2 females and 2 males per condition) from the AAV9-Control, AAV9-Low-TBX5-PHG and AAV9-TBX5-PHG. One group transduced with AAV9-hTNNT2-TBX5 (AAV9-TBX5) was added to enrich for the number of nuclei expressing TBX5 at higher levels (**Figure 2a**). Western blot analyses of isolated ventricular tissue validated increasing levels of TBX5 between the AAV9-Control, AAV9-Low-TBX5-PHG and AAV9-TBX5-PHG. AAV9-TBX5 vector expressed TBX5 at average levels comparable to those of AAV9-TBX5-PHG (**Figure 2b-c**). UMAP embedding demonstrated separate populations of nuclei of CM, endothelial cells, fibroblasts and immune cells (**Figure 2d-f, Supplementary Figure 1a-b**). We observed a decreasing fraction of endothelial nuclei and an increasing fraction of CM nuclei with increasing TBX5 dosage (**Figure 2e-f**). When comparing transcriptomes of AAV9-TBX5-PHG and AAV9-TBX5-derived nuclei to Control nuclei, respectively, we identified the most substantial transcriptional differences in CMs and endothelial cells (**Supplementary Figure 1c-f**). However, other non-CM cell types did not exhibit clear separation of graph-based clusters driven by the experimental treatment, suggesting limited sensitivity of intercellular (non-cell autonomous) signaling to TBX5 dosage in CMs (**Supplementary Figure 1g-l**).

**Figure 2.**
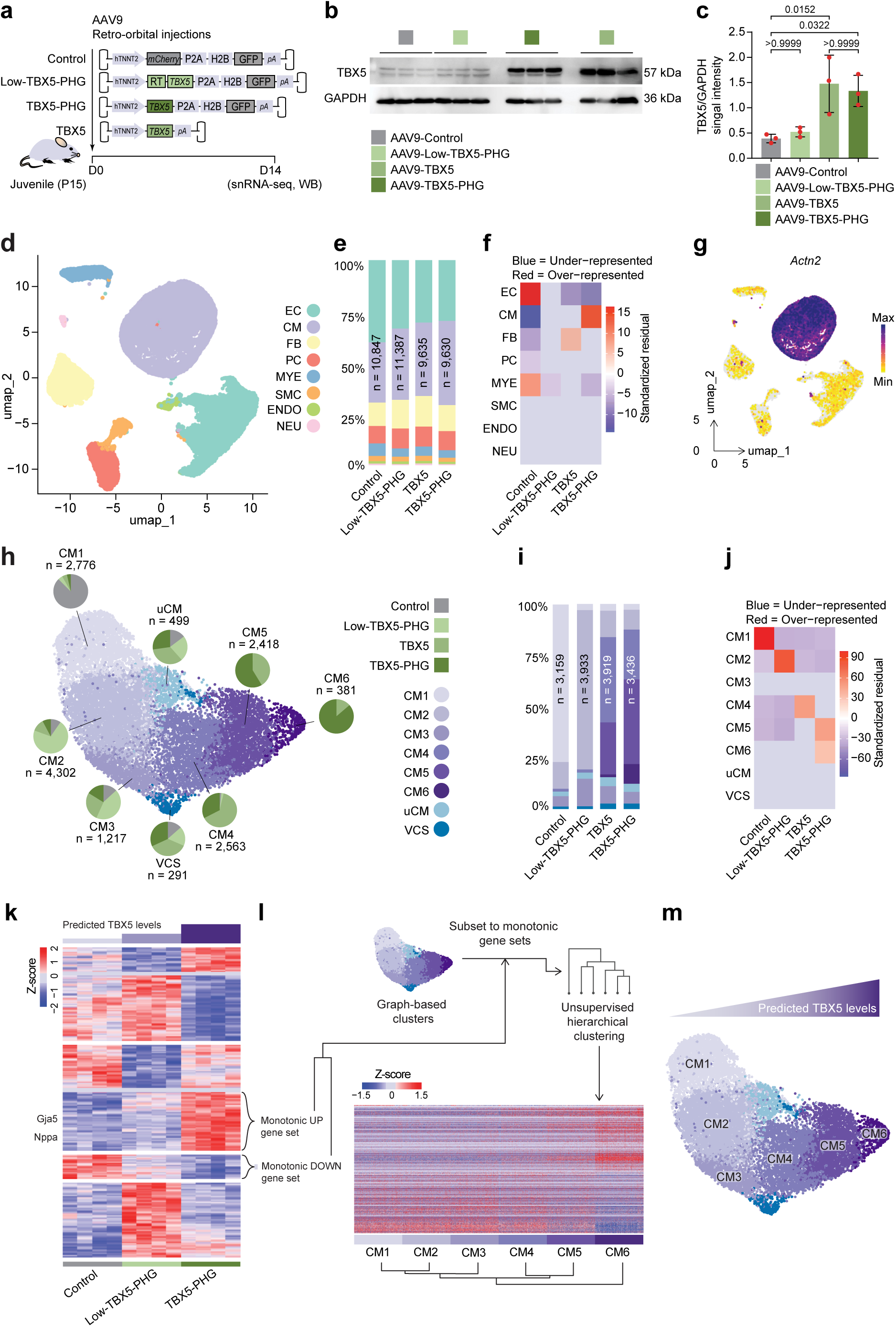
Single nucleus RNA-sequencing reveals CM-specific transcriptional responses to TBX5 dosage. (**a**) Schematic of experimental strategy for protein quantification and snRNA-sequencing, (**b**) Western blot of TBX5 protein levels from isolated ventricular mouse heart tissue, (**c**) Quantification of western blot showing TBX5 levels normalized by GAPDH (n=3), (**d**) UMAP embedding of all detected cell types in the snRNA-sequencing, performed on 4 pooled mouse hearts per condition, (**e**) Proportions of cell types per condition, (**f**) Chi-square test of cell type proportions per condition, (**g**) Feature plot showing *Actn2* expression on the UMAP of all detected cell types, (**h**) UMAP of reclustered CM showing CM1-CM6, uCM (unaffected CMs), and VCS (Ventricular conduction system) clusters. Pie charts indicate the proportion of each cluster contributed by each viral treatment condition, (**i**) Proportions of CM clusters per viral treatment condition, (**j**) Chi-square test of CM cluster proportions per condition, (**k**) Heatmap showing cluster analysis of DEGs in AAV9-Control, AAV9-Low-TBX5-PHG and AAV9-TBX5-PHG-treated mice (n=4), (**l**) (upper) Schematic showing the hierarchical clustering pipeline, (lower) Heatmap showing hierarchically clustered TBX5 dose-responsive genes (rows) and nuclei ordered by cluster (columns), (**m**) UMAP of CM clusters with indication of predicted TBX5 dosage gradient.

The CM population was reclustered and separated into eight main clusters (**Figure 2h-j, Supplementary Figure 1c-d**). Both clusters uCM (unaffected CMs) and ventricular conduction system (VCS), identified by commonly used VCS marker genes such as *Cntn2, Gja5, Scn10a*, *Etv1* and *Slit2,* contained comparable proportions of nuclei from all treatment conditions, suggesting they may be derived from the fraction of CMs that were not transduced with AAV expression vectors. Clusters CM1-CM6, on the other hand, showed marked shifts in composition, with one or more treatment groups disproportionately contributing to each cluster (**Figure 2h-j**). Chi-square analysis indicated that AAV9-Control nuclei were significantly overrepresented in CM1; AAV9-Low-TBX5-PHG in CM2 (and somewhat in CM3); AAV9-TBX5 in CM4; and AAV9-TBX5-PHG in both CM5 and CM6 (**Figure 2j**). These results suggest that CM clusters reflect distinct transcriptional states along a TBX5 dose-response continuum.

We next investigated whether transcriptional differences among CM clusters is indeed driven by varying levels of TBX5 activity. *TBX5* transcripts were poorly detected in the snRNA-seq data, and the inclusion of the uORF reduces TBX5 dosage post-transcriptionally, thus uncoupling mRNA and protein levels. Therefore, to estimate TBX5 activity levels, we compared bulk RNA-sequencing data of ventricular samples from three TBX5 dosage conditions (AAV9-Control, AAV9-Low-TBX5-PHG, and AAV9-TBX5-PHG). From this analysis, we identified target gene sets exhibiting strictly monotonic expression changes (either up- or downregulated) with increasing TBX5 dosage (**Figure 2k**). The transcriptional response of these two gene sets were subsequently used as proxies for TBX5 dosage. To assess the relationship between single-nucleus cardiomyocyte clusters (clusters CM1-CM6) and inferred TBX5 dosage, we randomly selected 500 nuclei per cluster and computed z-scored expression values for the monotonic gene sets. Hierarchical clustering was then applied to the normalized expression profiles while restricting inter-cluster mixing, such that hierarchical clustering determined the ordering of clusters rather than individual nuclei (**Figure 2l**, upper). This unsupervised analysis arranged the clusters (CM1-CM6) consistent with the progression observed along the UMAP trajectory (**Figure 2l**, lower). The resulting heatmap demonstrated that genes upregulated with higher TBX5 dosage in bulk RNA-seq showed progressively increased expression across the inferred ordering CM1→CM6, whereas genes downregulated with higher TBX5 dosage showed the opposite pattern (**Figure 2l**, lower). These results indicate that the six CM clusters likely represent successive transcriptional states along a predicted continuum of TBX5 dosages rather than representing discrete cell types (**Figure 2m**).

To analyze gene expression dynamics along the TBX5 dosage gradient, we applied Slingshot pseudotime analysis to CM1-CM6 (**Figure 3a**, upper). The expression patterns of TBX5 target genes *Nppa*, *Sbk2* and *Myl2* ^26,36^ plotted in the UMAP embedding were consistent with the predicted TBX5 dosage gradient (**Figure 3a**, lower). To further estimate the TBX5 levels across the TBX5 dosage gradient, we projected the normalized expression levels of TBX5-responsive genes *Nppa*, *Gja5*, *Plekha7*, *Parm1*, *Casq1*, and *Sbk2* in E16.5 ventricular trabecular and compact CMs ^68^ onto the pseudotime gradient. Target gene levels (e.g. *Nppa*) in E16.5 trabecular cells correspond roughly to the two-thirds point of the TBX5-dosage gradient. This estimate is in line with the dosage range measured in nuclei (**Figure 3b**).

**Figure 3.**
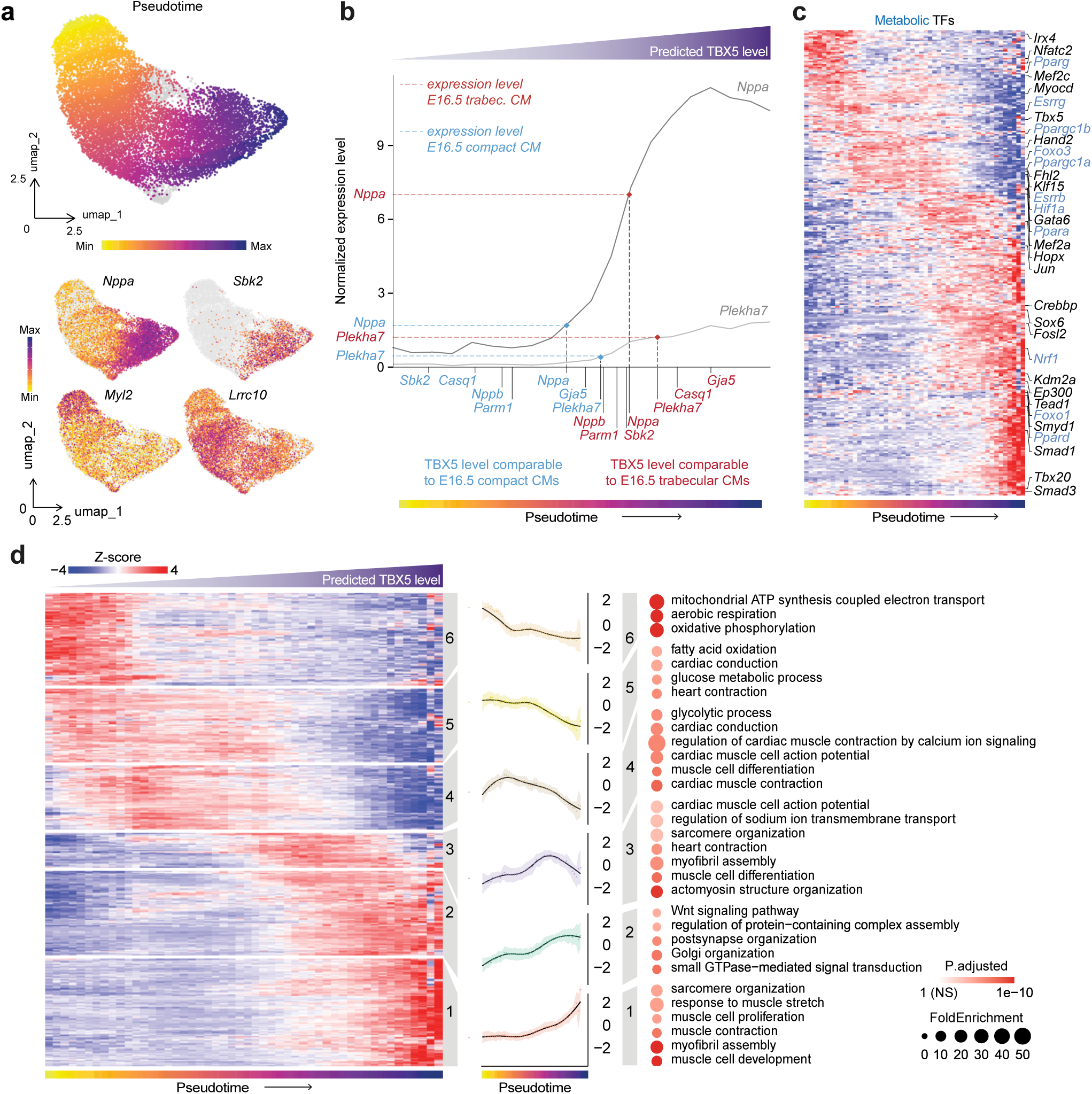
Pseudotime trajectory analysis reveals complex, non-monotonic transcriptional responses to TBX5 dosage. (**a**) UMAP of CM, coloring indicating Slingshot-calculated pseudotime (arbitrary units) (upper), (lower) feature plots showing *Nppa*, *Sbk2, Myl2* and *Lrrc10* expression, (**b**) Normalized expression of selected TBX5-responsive genes (*Nppa, Nppb, Gja5, Sbk2, Parm1, Casq1, Plekha7*) along the pseudotime trajectory of CM from TBX5 dosage snRNA-seq experiments, with dashed lines indicating expression levels in E16.5 fetal mouse trabecular and compact cardiomyocytes (data from GEO GSE132658), genes were normalized using the same set of reference genes used in Figure 1a, (**c**) Heatmap showing expression of all transcription factors across the 50 pseudotime bins, with selected transcription factors highlighted, (**d**) Heatmap showing the top 2,000 genes most strongly correlated with pseudotime, hierarchically clustered and cut into 6 gene clusters. Mid panels show expression profiles (average z-score per cluster), and right panel shows dot plot showing top relevant GO Biological Process (GO BP) terms for each gene cluster.

We then analyzed expression patterns of all TF genes found to be significantly correlated with pseudotime, and found diverse patterns throughout the trajectory, including metabolic regulators and several key cardiac TFs (**Figure 3c**). For example, at low dosages, high initial levels of *Pparγ*, a regulator of lipid metabolism and inflammatory signaling ^69,70^, were observed to decrease rapidly. This was followed at low dosages by the transient upregulation of *Ppargc1a* and *Ppargc1b*, coactivators essential for mitochondrial biogenesis and oxidative metabolism ^71^, alongside *Fhl2* and *Klf15*. *Fhl2* is a cardiac-specific scaffold protein known to modulate hypertrophic signaling, while *Klf15* serves as a critical inhibitor of pathological remodeling and fibrosis ^72,73^. At the highest dosages, the transcriptional response was characterized by the induction of *Tead1* and *Tbx20*. *Tead1* is a Hippo pathway effector required for CM survival and mitochondrial function, whereas *Tbx20* is a developmental factor whose postnatal overexpression is associated with fetal-like proliferative characteristics and myocardial repair ^70,72–76^. Collectively, these data indicate the cardiac TF network composition that governs functional and structural CM properties is highly sensitive to TBX5 dosage.

We performed GAM-based trajectory analysis to identify genes with significant expression dynamics along pseudotime (false discovery rate < 0.05), and selected the top 2000 genes with the strongest correlation across the 50 pseudotime bins for hierarchical clustering. We distinguished six gene clusters with their own temporal profiles (**Figure 3d**). Genes in cluster 6, expressed at low TBX5 levels and downregulated rapidly as TBX5 increases, are enriched for “oxidative phosphorylation”, thus suggesting that there is an initial and rapid suppression of mitochondrial respiratory activity in response to any dose of exogenous TBX5. Cluster 5 genes are enriched for “fatty acid oxidation” and “glycolysis”, implying that as TBX5 dosage increases, fatty acid oxidation is maintained while simultaneously a switch occurs to glycolysis for energy production. Cluster 4 genes show non-monotonic responses (for example *Lrrc10*; **Figure 3a, lower panel**) and are enriched for “cardiac conduction” and “differentiation” as well as “glycolysis”, suggesting moderate TBX5 levels support the emergence of some aspects of specialized CM functions. Cluster 3 is enriched for “sarcomere organization”, “heart contraction”, “myofibril assembly”, and “cardiac muscle action potential“; therefore, it appears that at intermediate TBX5 levels, the contractile apparatus may be enforced. Cluster 2 and cluster 1 are both upregulated at the highest levels of TBX5. However, while Cluster 2 remains relatively constant, cluster 1 genes continue to increase expression. Cluster 2 is enriched for Wnt signaling, Golgi organization, and small GTPase-mediated signal transduction. Cluster 1, enriched for “sarcomere organization”, “muscle cell proliferation”, and “muscle cell development”, suggests a dedifferentiated state with signs of cellular stress. Overall, our results demonstrate that TBX5 dosages lead to non-linear and non-monotonic transcriptional responses characterized by initial suppression of mitochondrial oxidation, subsequent upregulation of both fatty acid oxidation, glycolytic energy metabolism and contractile function at intermediate TBX5 levels, and dedifferentiation and stress at high TBX5 levels. These data reveal that different TBX5 dosages activate transcriptionally distinct cellular states.

### TBX5 confers conduction system-like identity to working cardiomyocytes

Several markers of the VCS were found to be upregulated at moderate-to-high dosages of TBX5, including well-characterized markers *Cntn2* and *Gja5* ^77,78^. Therefore, we aimed to investigate the degree to which a VCS-like identity was induced by these dosages of TBX5. *Cntn2* is typically expressed selectively in VCS cells after birth. When visualizing *Cntn2* expression in only CM nuclei of AAV9-Control or of AAV9-TBX5 transduced mice, respectively, we observed that in the AAV9-Control group, *Cntn2* is expressed in the VCS cluster whereas AAV9-TBX5 expresses *Cntn2* in both the VCS cluster as well as CM4 and CM5 (**Figure 4a**). To assess differential gene expression associated with the degree of VCS identity, we compared VCS versus CM1 and CM5 versus CM1, and plotted the corresponding log2 fold changes (L2FC) for both contrasts (**Figure 4b**). The strong diagonal correlation indicates that many of the genes defining VCS are upregulated in CM5, suggesting CM5 partially adopted a VCS state. Consistently, GJA5 is expressed throughout the ventricular wall in the AAV9-TBX5, where in the AAV9-Control ventricular free wall the signal is limited to Purkinje fibers of the VCS (**Figure 4d**). However, some VCS genes either do not show a change in their expression levels in response to TBX5 or show decreased levels. For instance, whereas VCS markers *Cntn2*, *Gja5* and *Scn10a* are induced by medium-high dosages of TBX5, *Etv1* and *Slit2* are strongly expressed in the VCS but are almost completely absent in CM4 and CM5, suggesting an incomplete VCS state induced by TBX5 (**Figure 4c**).

**Figure 4.**
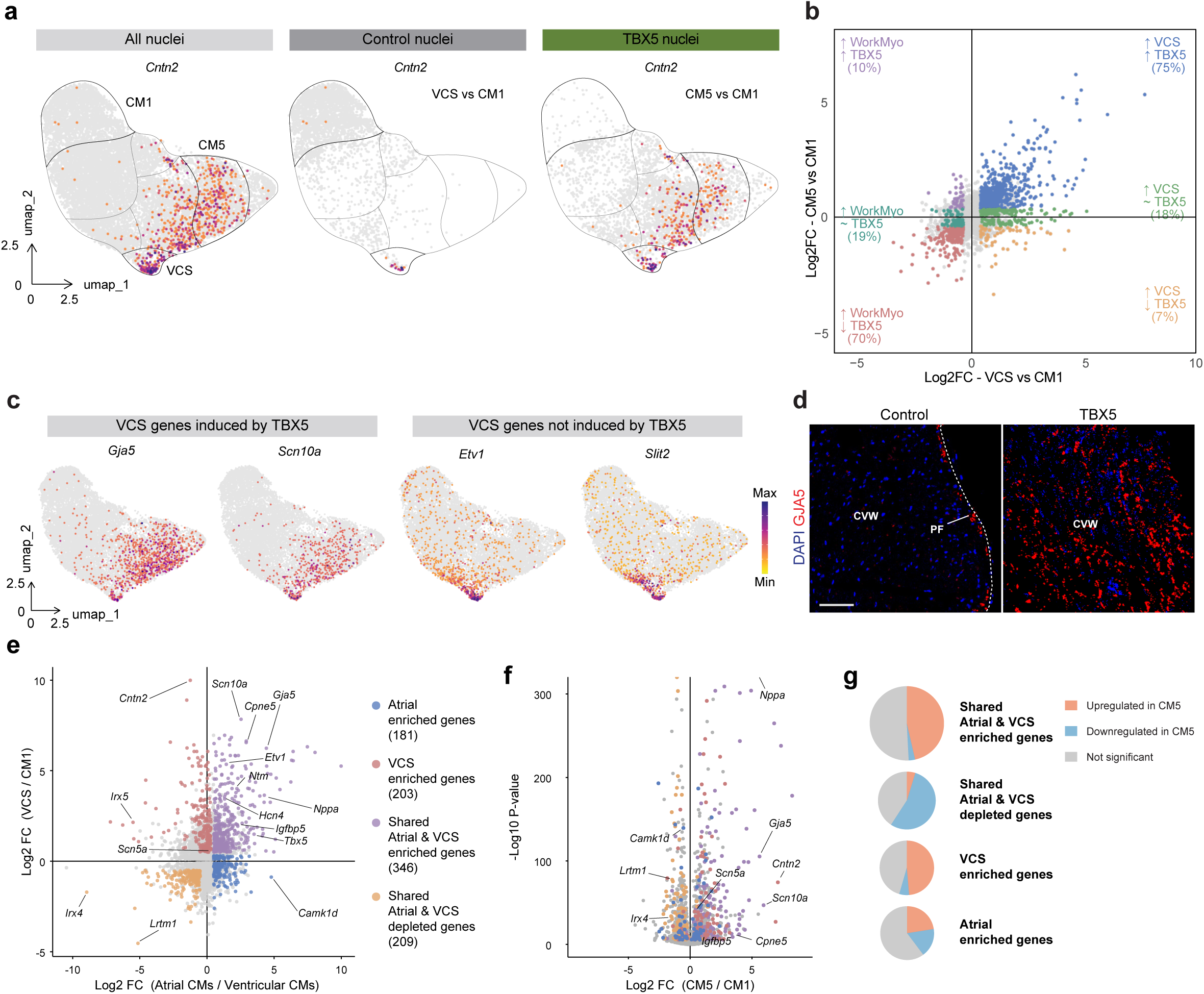
TBX5 overexpression induces partial ventricular conduction system identity. (**a**) Feature plots showing *Cntn2* expression across all cardiomyocytes (left), AAV9-Control nuclei (middle), and AAV9-TBX5 nuclei (right), (**b**) Scatterplot comparing Log2 fold change from differential expression analysis of CM5 versus CM1 (y-axis) and VCS versus CM1 (x-axis), (**c**) Feature plots showing expression of *Gja5, Scn10a, Etv1,* and *Slit2*, (**d**) Immunofluorescence microscopy images showing DAPI (blue) and GJA5 (red) in left compact ventricular wall (CVW) sections from AAV9-Control and AAV9-TBX5-treated mice (scale bar: 25 μm), (**e**) Scatterplot comparing log₂ fold changes (Log₂FC) of atrial vs. ventricular cardiomyocyte markers (GSE215021) against Log₂FC of VCS vs. CM1 markers (our data). Genes categorized by expression pattern: Shared Atrial and VCS enriched genes, Shared Atrial and VCS depleted genes, Atrial enriched genes, and VCS enriched genes, (**f**) Volcano plot (CM5 vs. CM1) highlighting the four gene categories from (**e**), (**g**) Pie charts indicating the fraction of each group of genes (from **e**) up- or downregulated by TBX5.

TBX5 was previously reported to maintain atrial identity, and to impose that on ventricular CMs ^36^. However, there is substantial overlap between atrial CM-enriched and VCS-enriched gene expression when compared to ventricular CMs (**Figure 4e-f**). To assess whether TBX5 drives ventricular cardiomyocytes toward an atrial or VCS-like state, we first defined gene sets enriched in atrial CMs and VCS relative to ventricular CMs (**Figure 4f**) ^62^. We analyzed the overlap in proportions of differentially expressed genes (DEGs). 346 genes were commonly upregulated in both atria and VCS, among which 46.5% were upregulated and 2.9% downregulated by TBX5 in ventricular CMs. Of the 209 genes commonly downregulated in atria and VCS, 54.5% were downregulated and 4.8% upregulated by TBX5 in ventricular CMs. Additionally, 203 genes were uniquely VCS enriched (48.8% upregulated and 5.9% downregulated by TBX5), and 181 were uniquely atrial enriched (22.7% upregulated and 17.1% downregulated by TBX5) (**Figure 4f-g**). Highly specific VCS markers *Cntn2, Scn10a* and *Gja5* were among the genes upregulated by TBX5 in the ventricular CMs (**Figure 4f**). These data indicate that TBX5 in the ventricles induces a state more similar to VCS than to atrial CM state.

### TBX5 transduction attenuates CM hypertrophy and transcriptional changes associated with *Nppa-Nppb* deficiency

We previously generated a mouse line in which the *Nppa*-*Nppb* gene cluster was deleted from the genome (*Nppa-Nppb^⁻/⁻^*) ^42^. These mice show a significant increase in heart size due to CM hypertrophy ^61^. Their RNA-seq data of ventricular tissue revealed modest transcriptional changes in *Nppa-Nppb^⁻/⁻^* mice, including downregulation of programs for “oxidative phosphorylation”, “regulation of cardiac muscle contraction” and “cardiac muscle cell action potential” (**Supplementary Figure 2a-b**, **Supplementary File, Tables S4-5**). Notably, ventricular *Tbx5* expression is significantly downregulated, as are a number of VCS marker genes, including *Scn10a* and *Hcn4*, indicative of a broader reduction in the VCS transcriptional program (**Supplementary Figure 2a**; **Supplementary File, Table S6**). These observations suggest that reduced TBX5 dosage may contribute to the transcriptional changes and hypertrophic state of the ventricles of *Nppa-Nppb^⁻/⁻^* mice. To test this possibility, we retro-orbitally delivered AAV9-Control or AAV9-TBX5-PHG to *Nppa-Nppb^⁻/⁻^* mice (**Figure 5a**). After 2 weeks, delivery and CM-specific GFP expression were confirmed (**Figure 5b**). The CM size distributions of the groups indicated a significant reduction in CM sizes in AAV9-TBX5-PHG hearts compared to Control hearts, even though average CM sizes only showed a trend towards reduced sizes (**Figure 5c-d**). PCA of bulk RNA-sequencing data of ventricular tissue showed separation between the Control and the AAV9-TBX5-PHG treated *Nppa-Nppb^⁻/⁻^* mice (**Figure 5e**). Genes upregulated by TBX5 in *Nppa-Nppb^⁻/⁻^* ventricles were associated with “regulation of cardiac muscle contraction”, “regulation of atrial cardiac muscle cell membrane depolarization”, and “bundle of His cell to Purkinje myocyte communication”. No terms were significantly associated with the downregulated genes (**Figure 5f**, **Supplementary Figure 2c, Supplementary File, Tables S7-9**).

**Figure 5.**
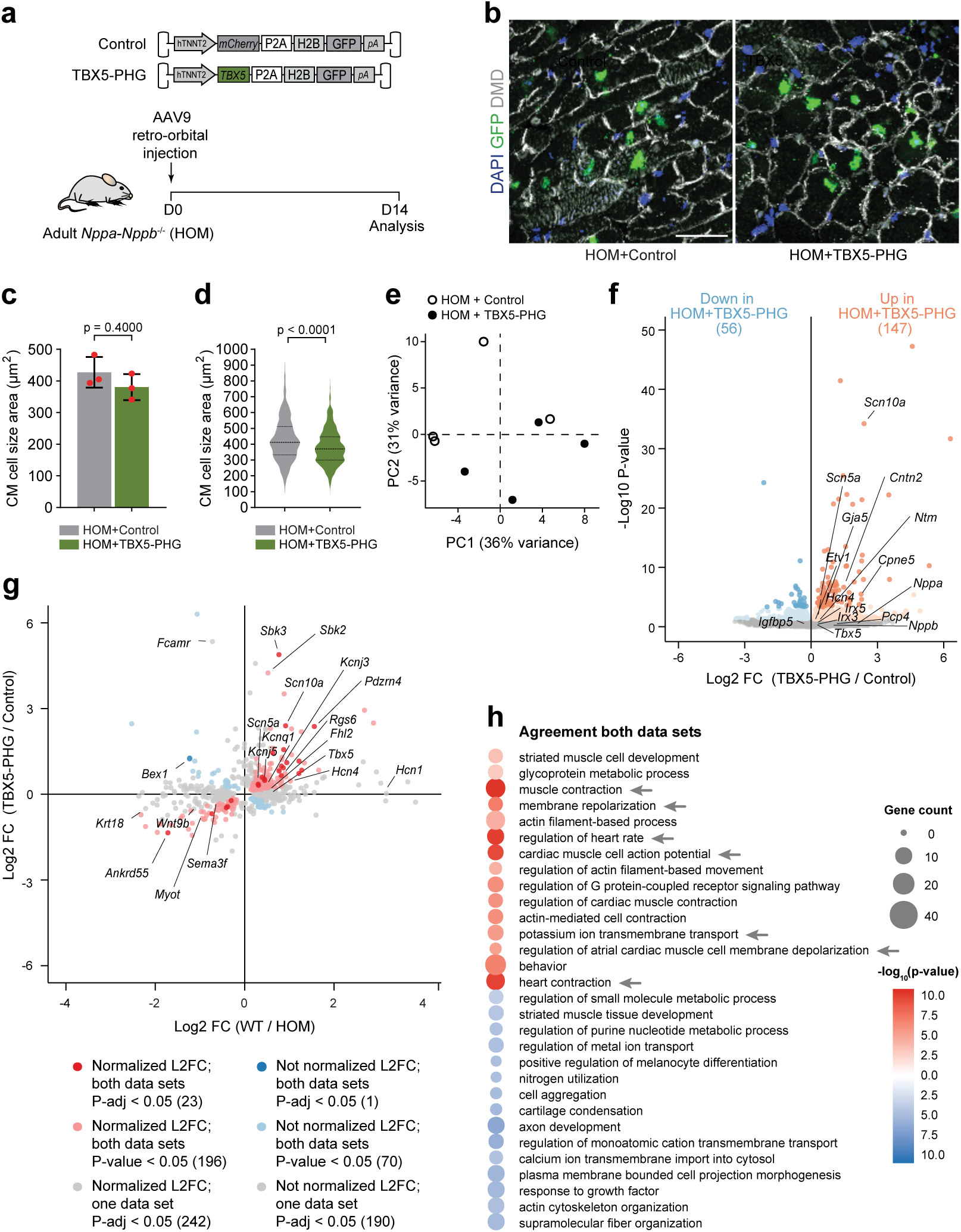
TBX5 partially restores the conduction system related transcriptomic profile observed in *Nppa-Nppb^⁻/⁻^* mice. (**a**) Schematic of *in vivo* experimental model in 8 week *Nppa-Nppb^⁻/⁻^* (HOM) mice, (**b**) Immunohistochemistry showing successful transduction of the AAV9-control and AAV9-TBX5-PHG as visualized by GFP (green) specifically in CM as marked by DMD (gray), and nuclei depicted in blue (scale bar: 25 μm), (**c**) Quantification of the cell size average (n=3), (**d**) Cell size area distribution (n=3) in the ventricles of *Nppa-Nppb^⁻/⁻^* (HOM) mice injected with, AAV9-Control or AAV9-TBX5-PHG (CM distribution area (60 CMs per section/over 3 sections per heart), (**e**) Principal Component Analysis (PCA) plot showing separation between *Nppa-Nppb^⁻/⁻^* (HOM) mice injected with either AAV9-control (n=4) or AAV9-TBX5-PHG (n=4), (**f**) Volcano plot showing 203 DEGs between *Nppa-Nppb^⁻/⁻^* (HOM) mice injected with AAV9-control (n=4) and AAV9-TBX5-PHG (n=4), (**g**) Scatterplot comparing gene expression changes in *Nppa-Nppb^⁻/⁻^* hearts (x-axis, log₂FC WT vs. HOM) with the transcriptional response to TBX5 administration (y-axis, log₂FC TBX5 vs. control). Each dot represents a single gene, dark red and dark blue points indicate genes significantly differentially expressed in both comparisons (P_adj_ < 0.05), light red and light blue points indicate nominal significance in both comparisons (P < 0.05), and gray points indicate genes significant in only one comparison. Selected ventricular conduction system markers are labeled, (**h**) Dot plot showing enriched biological process Gene Ontology (GO) terms among genes normalized by TBX5 overexpression in *Nppa-Nppb^⁻/⁻^* (HOM) hearts (15 terms). Dot color reflects significance (red, upregulated; blue, downregulated), and dot size indicates the number of genes from the input set annotated to each term. Comparisons between two groups were analyzed using the Mann-Whitney U test. The distribution of CM size was analyzed using the Kolmogorov-Smirnov test, where 180 cells per heart (n=6 per group) were pooled together.

To assess the extent to which TBX5 delivery restored gene expression toward wild-type levels in *Nppa-Nppb^⁻/⁻^* hearts, we compared the magnitude and direction of differential expression for each gene between the knockout comparison (*Nppa-Nppb^+/+^* (WT) vs *Nppa-Nppb^⁻/⁻^*) and the TBX5 treatment comparison (AAV9-TBX5-PHG vs AAV9-Control in *Nppa-Nppb^⁻/⁻^*) (**Figure 5g**). In this plot, positive x-axis values indicate lower expression in HOM hearts relative to WT, and positive y-axis values indicate increased expression following TBX5 treatment. Genes showing opposite-direction changes between the two comparisons, indicating that TBX5 shifted expression toward the WT state, were classified as restored. Among the 24 genes that were significantly differentially expressed in both datasets (P_adj_ < 0.05), 23 were restored. When the analysis was expanded to include all genes showing nominal significance in both comparisons (P < 0.05), 75.5% (219/290) showed restoration, whereas 24.5% (71/290) changed in the same direction in both comparisons and were therefore not restored. Gene Ontology enrichment analysis of the 219 restored genes revealed significant enrichment for processes related to cardiac electrophysiology and contractile function (**Figure 5h**). Enriched terms included regulation of “atrial cardiac muscle cell membrane depolarization”, “cardiac muscle cell action potential”, “regulation of heart rate”, and “muscle contraction”, indicating that TBX5 overexpression preferentially restored expression of genes associated with electrical activity and functional properties of cardiomyocytes (**Supplementary File, tables S10-11**). Notably, genes associated with the VCS, including *Gja5, Scn10a, Scn5a and Kcnj3* were enriched among the normalized genes. These data suggest that reduced expression of *Tbx5* contributes to the transcriptomic changes and CM hypertrophic state in *Nppa-Nppb^⁻/⁻^* ventricles.

### Ventricular phenotypic responses to TBX5 dosage increases

To investigate the phenotypic responses of the ventricles to increasing TBX5 dosages, we transduced mice with AAV9-Control, AAV9-Low-TBX5-PHG and AAV9-TBX5-PHG (**Figure 6a**). The transduction efficiency was quantified based on the GFP^+^/PCM1^+^ signal (**Supplementary Figure 3a**) and reached consistently 80% for the AAV9-Control and AAV9-TBX5-PHG constructs. AAV9-Low-TBX5-PHG reached only 50% transduction efficiency, which may result from TBX5 (and H2B-GFP) in a fraction of the nuclei being expressed at levels below the detection limit of our assay (**Figure 6b**). Based on the TBX5 dosages observed in ventricles transduced with these vectors (**Figure 6b**) and the cluster assignment in the snRNA-sequencing analysis (**Figure 2g**), we predicted the average TBX5 dosage in Control ventricular CMs represent the dosage in cluster CM1, in AAV9-Low-TBX5-PHG the dosage in CM2-3, and in AAV9-TBX5-PHG the dosage in CM4-6. To validate the TBX5 dosage-transcriptional response prediction, we performed RNA-seq analysis of ventricular tissue of these transduced mice. Principal component analysis (PCA) showed that the different TBX5 dosages accounted for the largest fraction of variation (**Figure 6c**). Comparison between data sets revealed 1127 DEGs between AAV9-Low-TBX5-PHG and AAV9-Control, and 1255 DEG between AAV9-TBX5-PHG and Control, whereas fewer genes (405) were differentially expressed between both TBX5 groups. The overall magnitude of change in expression levels was higher for AAV9-TBX5-PHG vs Control compared to Low-TBX5-PHG vs Control (**Figure 6d**). Cluster analysis revealed gene sets that responded either monotonically or non-monotonically to TBX5 dosage (**Figure 6e**). GO term analysis of the clusters validated the responses of gene programs associated specifically with AAV9-Control, AAV9-Low-TBX5-PHG or AAV9-TBX5-PHG observed in the snRNA-seq analysis (**Figure 6f**). Thus, AAV9-Low-TBX5-PHG ventricles showed increased expression of genes associated with lipid metabolism, fatty acid oxidation and the tricarboxylic acid cycle (e.g. *Ech1*, *Cpt2, Idh2*), sarcomere organization, calcium handling and contractility (*Lrrc10*, *Mybpc3*) ^79^, whereas these were depleted in AAV9-TBX5-PHG (**Supplementary File, Table S2**). At high TBX5 dosage (AAV9-TBX5-PHG), terms associated with gene programs involved in (de)differentiation (*Erbb2, Nrg1*) and cardiac conduction (*Gja5)* were enriched. In general, VCS markers (*Cntn2, Scn10a*) were upregulated most strongly by the high TBX5 dosage (AAV9-TBX5-PHG) (**Supplementary File, Table S3**).

**Figure 6.**
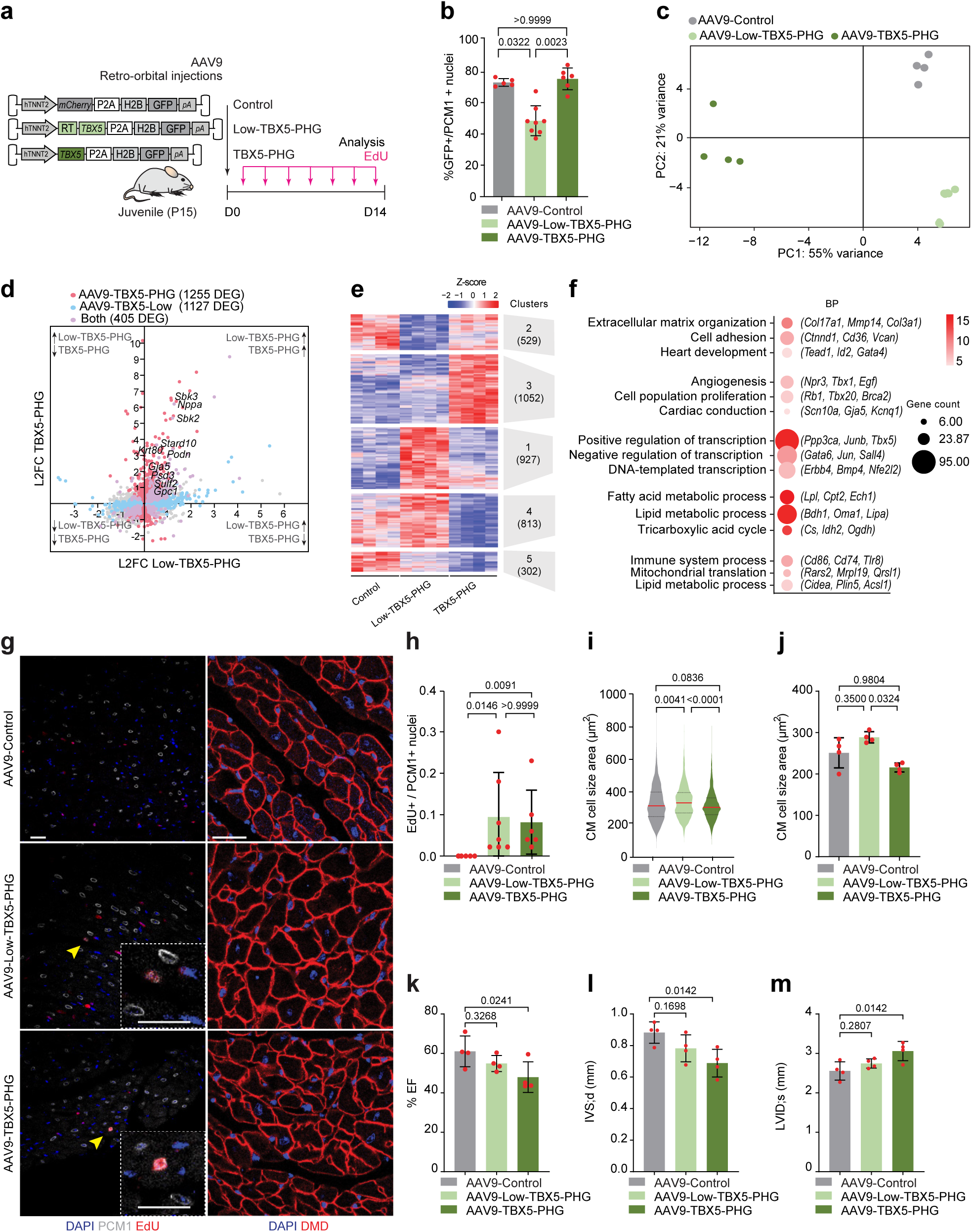
Two doses of AAV9-TBX5 result in opposite transcriptional and ventricular states. (**a**) Schematic of *in vivo* experimental model in P15 mice, (**b**) Quantification of GFP^+^/PCM1^+^ nuclei (Control; n=5, Low-TBX5-PHG; n=8, TBX5-PHG; n=6), (**c**) Principal component analysis (PCA plot) of bulk RNA sequencing for AAV9-Control, AAV9-Low-TBX5-PHG and AAV9-TBX5-PHG-treated mice (n=4), (**d**) Scatter plot comparing the log₂FC between AAV9-Low-TBX5-PHG and AAV9-TBX5-PHG mice. Red, blue and purple dots indicate genes that were differentially expressed (P_adj_<0.01) only in AAV9-TBX5-PHG, only in AAV9-Low-TBX5-PHG or both, respectively (n=4), (**e**) Heatmap showing cluster analysis of DEGs in AAV9-Control, AAV9-Low-TBX5-PHG and AAV9-TBX5-PHG-treated mice (n=4; same data as Figure 2k), (**f**) Gene ontology (GO analysis) showing shared biological processes in mice injected with AAV9-Low-TBX5-PHG and AAV9-TBX5-PHG, (**g**) Representative immunofluorescence staining of PCM1 (gray), GFP (green), EdU (red), and DAPI (blue) (left), and DMD (red) and DAPI (blue) (right) of ventricles from AAV9-Control, AAV9-Low-TBX5-PHG, and AAV9-TBX5-PHG injected mice. Scale bar: 25 μm, (**h**) Quantification of EdU^+^/PCM1^+^ nuclei (Control; n=5, Low-TBX5-PHG; n=7, TBX5-PHG; n=6), (**i**) Quantification of the cell size distribution (n=4), (**j**) Average of cell size area (n=4) in the ventricles of AAV9-Control, AAV9-Low-TBX5-PHG and AAV9-TBX5-PHG-treated mice, (**k-m**) Echocardiography measurements of (**k**) ejection fraction (EF), (**l**) interventricular septum thickness (IVS;d) and (**m**) left ventricular internal diameter (LVID;s) in the ventricles of AAV9-Control, AAV9-Low-TBX5-PHG and AAV9-TBX5-PHG-treated mice (n=4). Data represented as mean ± SD. Each dot represents a biological replicate (n). Comparison between more groups were analyzed by Kruskal-Wallis. The distribution of CM size was analyzed using the Kolmogorov-Smirnov test, where 180 cells per heart (n=4 per group) were pooled together.

We next analyzed phenotypic responses of the ventricles of transduced mice. HW/BW and HW/TL ratios did not differ between control, AAV9-Low-TBX5-PHG and AAV9-TBX5-PHG transduced hearts (**Supplementary Figure 3c-d**). There was a significant increase in EdU^+/^PCM1^+^ nuclei in both AAV9-Low-TBX5-PHG and AAV9-TBX5-PHG compared to the control mice (**Figure 6g**, left column**-h**). Interestingly, quantification of the DMD^+^/DAPI^+^ cross-sectional CM area revealed a consistent pattern, with AAV9-Low-TBX5-PHG caused increasing CM size (**Figure 6g**, right column; **Figure 6i-j**), whereas AAV9-TBX5-PHG exhibited reduced CM size, **(Figure 6i-j**). Echocardiography analysis 2 weeks after injection showed no systolic (s) or diastolic (d) differences between AAV9-Control and AAV9-Low-TBX5-PHG injected mice, and the left ventricular (LV) mass was unaffected. However, AAV9-TBX5-PHG-transduced mice showed reduced ejection fraction and fraction shortening (EF, FS), and interventricular septum thickness (IVS;d), and increased left ventricular internal diameter (LVID;s, LVID;d) and left ventricular volume (LV vol;d, LV vol;s). Other parameters remained unchanged upon AAV9-TBX5-PHG delivery (**Figure 6k-m**, **Supplementary Figure 3d**).

To assess the functional impact of OXPHOS and FAO gene downregulation of OXPHOS and FAO genes in hearts expressing relatively high levels of TBX5, we transduced mice with AAV9-GFP (Control) or AAV9-TBX5 (**Figure 7a**). High cardiac-specific transduction efficiency was confirmed (**Supplementary Figure 4a-c**), with no morphological or fibrotic differences between the two groups (**Supplementary Figure 4d-h**). Activation of the cell cycle and suppression of the OXPHOS and FAO programs (**Supplementary Figure 4i-l**), and CM size reduction and CM cell cycle re-entry by AAV9-TBX5 were validated (**Figure 7b-d, Supplementary Figure 4m-s**). We measured mitochondrial respiration in ventricles of mice transduced with AAV9-GFP or AAV9-TBX5. A significant decrease in complex I (CI), as well as the total OXPHOS and the uncoupled maximal respiration rates was observed in AAV9-TBX5 ventricles. Complex II (CII) was not significantly changed but showed a tendency to be downregulated (**Figure 7e-f**). While the overall mitochondrial respiratory capacity was diminished, NADH-linked/OXPHOS, succinate-linked/uncoupled and ND1/HK2, 16S/HK2 ratios were not significantly different between AAV9-GFP and AAV9-TBX5 ventricles (**Figure 7g-h**, **Supplementary Figure 4t-u**). These results indicate preserved mitochondrial integrity and function. Similarly, TBX5 overexpression decreased FAO respiratory capacity (**Figure 7f**). Together, these data suggest that postnatal expression of TBX5 can promote hallmarks of CM dedifferentiation, including cell cycle re-entry, CM size reduction, and reduced mitochondrial respiration and FAO.

**Figure 7.**
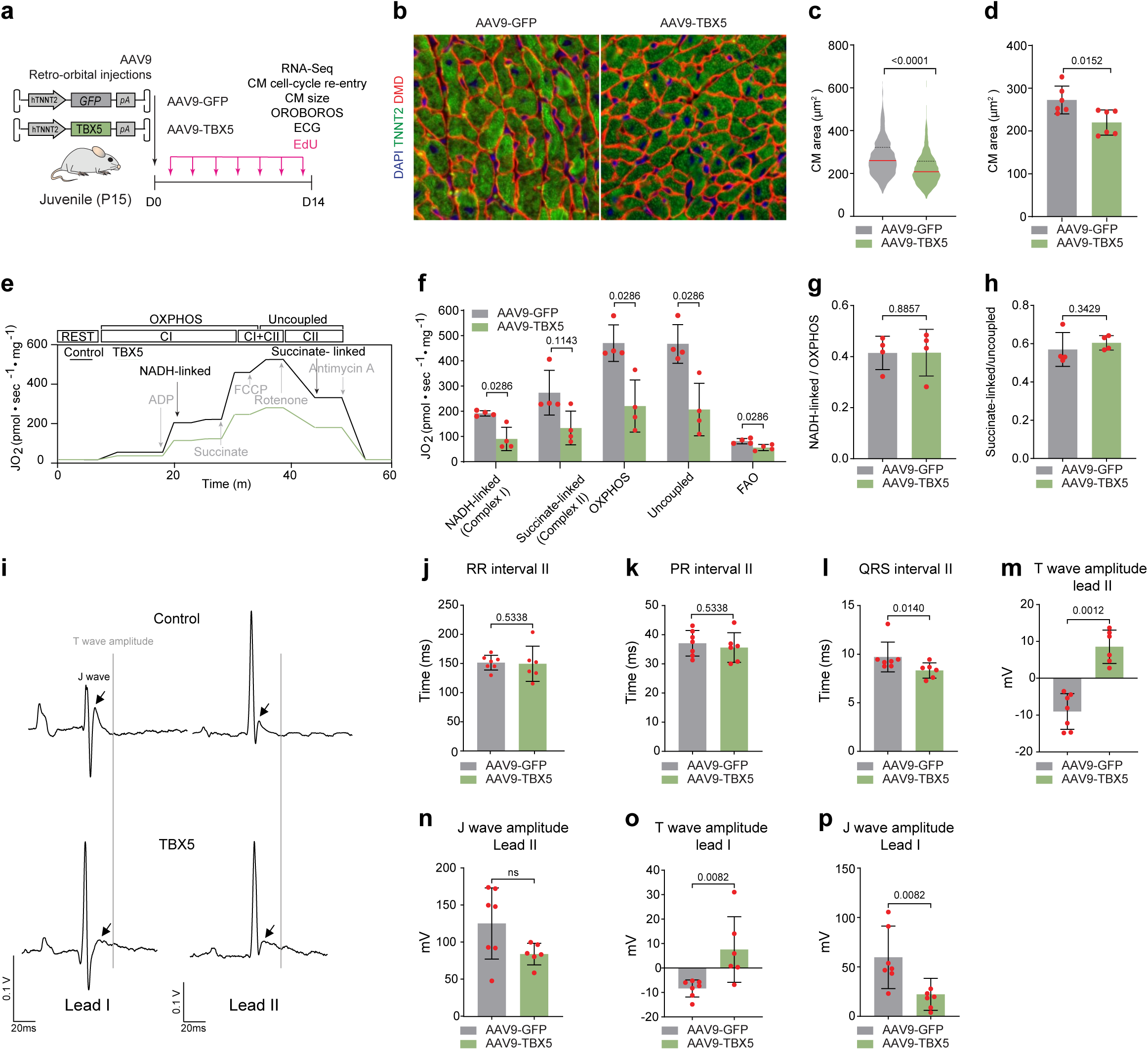
CM-specific TBX5 delivery profoundly changes the transcriptional profile of the postnatal ventricles. (**a**) Schematic of *in vivo* experimental model in P15 mice, (**b**) Immunostaining for TNNT2 (green), DMD (red) and DAPI (blue) in the ventricles of AAV9-Control and AAV9-TBX5-treated mice (scale bar: 25 μm), (**c**) Quantification of CM distribution area (60 CMs per section/over 3 sections per heart), (**d**) Quantification of CM cell size average area (n=6), (**e**) Schematic representation of the Oroboros experimental design, (**f**) Flux of O_2_ consumption on isolated alive tissue. The graph depicts the quantification of complex I, complex II, OXPHOS, uncoupled and FAO respiratory capacity, respectively (n=4), (**g**) NADH-linked/OXPHOS ratio, (**h**) succinate-linked/uncoupled ratio quantifying mitochondrial respiration efficacy (n=4), (**i**) Examples of surface ECGs in AAV9-GFP (Control) and AAV9-TBX5 mice, with T wave and J wave amplitudes depicted, ((**j**) RR interval, (**k**) PR interval, and (**l**) QRS duration (lead II), (**m-p**) T wave amplitude and J wave amplitude measurements for lead I and lead II in AAV9-Control (n=7) and AAV9-TBX5-treated mice (n=6). Data represented as mean ± SD. Each dot represents a biological replicate (n). Comparisons between two groups were analyzed using the Mann-Whitney U test. Comparison between more groups were analyzed by Kruskal-Wallis. The distribution of CM size was analyzed using the Kolmogorov-Smirnov test, where 180 cells per heart (n=6 per group) were pooled together.

The snRNA-seq analysis indicated the CMs in clusters CM4-5 (in which AAV9-TBX5 was enriched) acquired a VCS-like state and was accompanied by overexpression of GJA5 in the ventricular myocardium. To assess whether this state causes electrophysiological alterations, we performed surface electrocardiography in mice transduced with either AAV9-GFP or AAV9-TBX5 (**Figure 7i**). RR and PR intervals were comparable between AAV9-GFP and AAV9-TBX5 mice whereas QRS duration was shortened, indicative of increased conduction velocity (**Figure 7j-l**) and ventricular repolarization altered by increasing T wave amplitude and decreasing J wave amplitude in AAV9-TBX5 mice (**Figure 7m-p**). These findings are consistent with the TBX5-induced VCS state in ventricular CMs. Namely upregulation of *Gja5*, *Cntn2*, and *Scn10a*, and downregulation of the calcium handling genes *Ryr2*, *Pln* and *Atp2a2* (**Supplementary File, Table S12**).

Collectively, these observations indicate that increasing TBX5 dosages lead to distinct functional changes in CMs, including reduced mitochondrial oxidative capacity and altered ventricular conduction properties, in line with the transcriptional state changes. Slightly increased TBX5 dosages promoting CM size and programs associated with a more mature state, and higher TBX5 dosages promoting CM dedifferentiation and reduced functionality.

## Discussion

Numerous studies have underscored the profound impact of transcription factor (TF) dosage on molecular and cellular states ^3–11^. However, few experimental studies have addressed how gradual variations in TF dosage across physiologically relevant ranges dictate the state of fully differentiated cells *in vivo*. In this study, we titrated the expression of the dosage-sensitive TF TBX5 across a physiologically plausible spectrum in postnatal ventricular CMs. Our results demonstrate that transcriptional and phenotypic responses to TBX5 are inherently non-linear and non-monotonic, characterized by striking shifts in TF network composition and gene programs governing CM properties such as metabolism and conductivity.

A critical premise of our model is that the experimental TBX5 dosages reflect physiological conditions. Our integrated analyses indicate that the transduced CMs span a dosage continuum from endogenous adult ventricular levels to levels exceeding those of the adult atrium, including intermediate levels observed in fetal trabecular myocardium and the postnatal ventricular conduction system (VCS). This model is therefore uniquely suited to investigate the dosage-phenotype relationships that underlie complex traits, disease risk, and more severe dysregulation ^19,38–40^.

Unsupervised clustering revealed that TBX5 dosage was the primary driver of transcriptional variation in CMs. This aligns with the known capacity of TBX5 to reprogram transcriptomes ^8^ and the high sensitivity of cardiac morphogenesis to TBX5 haploinsufficiency ^32,80^. Our snRNA-seq analysis of fully differentiated CMs *in vivo* revealed distinct dose-dependent states: at the lower end of the titration range, CMs acquired a state associated with enhanced lipid metabolism, sarcomere organization, and conductivity, which were phenotypically characterized by CM cell cycle re-entry, increase in CM size, and preserved cardiac function. Conversely, higher dosages induced downregulation of genes involved in oxidative phosphorylation (OXPHOS), fatty acid oxidation (FAO), calcium handling, and contractility, while simultaneously upregulating fetal- and proliferation-associated programs. These transcriptional shifts manifested phenotypically as cell cycle re-entry, reduced CM size, diminished ejection fraction, and impaired metabolic capacity. These observations indicate that non-linear, and specifically non-monotonic, responses dominate the landscape of mature CMs. While many target genes remain stable under minor fluctuations, others are exquisitely sensitive to subtle variations ^10^. Our findings align with genetic evidence suggesting that subtle TBX5 changes (<50% variation) drive trait variation, such as cardiac conductivity and atrial fibrillation risk ^19,21,41^. These effects are qualitatively and quantitatively distinct from the severe developmental disorders caused by major dosage deviations (≥50%), such as those seen in classical Holt-Oram Syndrome ^25–28,38^.

We further identified a large cohort of TF genes sensitive to TBX5 dosage, including key metabolic regulators such as *Pparγ* and *Ppargc1b*, which drive FAO in mature CMs ^70,76^. This highlights that TFs operate within complex, evolved regulatory networks that maintain homeostasis by coordinating shifts in network composition in response to dosage cues ^3,8,10^. Recent evidence has specifically implicated TBX5 in CM metabolism ^81,82^. For instance, TBX5 expression in iPSC-derived CMs promotes metabolic and structural maturity ^83^. Our data refine this understanding by uncovering that modest increases in ventricular TBX5 support a metabolically mature state, whereas increases beyond a particular dosage induce a fetal-like dedifferentiation characterized by reduced respiratory capacity.

TBX5 dosage may serve as a mechanism for phenotypic adaptation. *TBX5* levels are reduced in patients with dilated cardiomyopathy and heart failure, as well as in mouse models of hypertrophy, including our *Nppa-Nppb*-deficient model ^61,81,84^. Interestingly, in mice, chronic hypoxia, which confers resistance to pressure overload, upregulates TBX5, as does myocardial recovery following ventricular assist device implantation in humans ^84^. By demonstrating that increasing TBX5 dosage in *Nppa-Nppb*-deficient mice partially rescues gene dysregulation and hypertrophy, we propose that TBX5 titration is an adaptive mechanism to cope with cardiac stress. This suggests that the elevated *TBX5* levels associated with atrial fibrillation ^19,21^ might also reflect a functional, albeit potentially maladaptive, compensatory response. Future studies into the causal relationships between TBX5 modulation and CM adaptation will be essential to further elucidate these homeostatic mechanisms.

Within a specific TBX5 dosage window, we observed that a substantial fraction of VCS-enriched genes was upregulated in ventricular working CMs. The VCS component in our model primarily comprises Purkinje fiber CMs, specifically marked by *Gja5* ^77^, *Cntn2* ^78^ and *Scn10a* ^85,86^. During development, VCS CMs diverge from fetal trabecular CMs, which express approximately threefold higher levels of *Tbx5* than compact CMs, a differential that is maintained in the postnatal heart. Remarkably, elevating TBX5 dosage in mature working CMs to levels approximating those of the VCS was sufficient to induce a VCS-like transcriptional state, suggesting considerable phenotypic plasticity within CM subtypes. However, certain key VCS genes, such as the TF *Etv1* ^87^, were not induced, indicating that while TBX5 is a potent driver, it is not sufficient to impose a full VCS identity on ventricular CMs. Furthermore, the *Nppa-Nppb*^⁻/⁻^ model, characterized by CM hypertrophy and reduced *Tbx5* expression, exhibited a downregulation of VCS markers. The fact that AAV9-TBX5 delivery rescued a large portion of these transcriptional changes (including *Gja5*, *Scn5a*, and *Hcn4*) positions these mice as a valuable model for studying the relationship between reduced TBX5 dosage and ventricular traits.

At the upper end of the dosage spectrum, high levels of TBX5 upregulated genes associated with Hippo and Neuregulin (*Erbb2*) signaling. The state of these high-TBX5 CMs resembles that of adult CMs that have dedifferentiated in response to *Nrg1*/*Erbb2* activation, Hippo pathway modulation, or metabolic and epigenetic remodeling to reacquire regenerative potential ^88–91^. A subpopulation of ventricular CMs was identified that transiently reactivates *Tbx5* upon injury; however, while these cells adopt a dedifferentiated state, they did not inherently stimulate regeneration ^92^. Our findings suggest that prolonged TBX5 overexpression may be detrimental to cardiac function. Whether temporal TBX5 modulation can enable diseased CMs to dedifferentiate, regenerate, and subsequently redifferentiate to provide protection against stress or ischemia ^93^ remains an open question.

In summary, modulating TBX5 dosage *in vivo* induces non-monotonic transcriptional and phenotypic states in cardiomyocytes, driven by a non-linear relationship in which lower levels enhance CM maturity while higher levels trigger CM dedifferentiation and deterioration of cardiac function. By defining this dose-dependent landscape, this study establishes a semi-quantitative framework for how subtle stoichiometric shifts in transcription factor activity dictate the divergence between physiological maturation and disease states.

## Author Contributions

A.E. Giovou, O.J. Mulleners, M.M. Gladka and V.M. Christoffels designed the experiments. A.E. Giovou, O.J. Mulleners, R. Caliandro, A.R. Boender, and V.A.J. Warnaar performed all experiments. A.E. Giovou, O.J Mulleners, V.A.J. Warnaar, M.R. Rivaud and V.M. Christoffels analyzed the data. M. Klerk and M. Giacca provided essential reagents including AAV constructs. A.E. Giovou, O.J Mulleners, M.M. Gladka and V.M. Christoffels wrote the manuscript. All authors revised the article.

## Funding

This work was supported by European Innovation Council Pathfinder Challenges Project 101115295 - Nav1.5-CARED and TRANSITION - Project 101099608 - TRACTION (to V.M.C. and G.J.J.B.); Prof. Dr. AF Moorman Fund of the Amsterdam University Fund (to B.J. and V.M.C.); Dutch Heart Foundation and Hartekind Foundation Grant CVON2019-002 OUTREACH (to V.M.C.); Health Holland PPP-Grant 2024 CURE-VT/VF (to G.J.J.B. and V.M.C.); Aspasia research programme financed by the Dutch Research Council (NWO) (015.021.029) (to M.M.G.); Dr. Dekker Senior Scientist Fellowship from the Dutch Heart Foundation (NHS2020T041 to M.M.G); M.R.R. was supported by the Dutch Heart Foundation and Dutch CardioVascular Alliance 01-002-2022-0118 EmbRACE (to V.M.C.).

## Conflict of interests

G.J.J.B. reports ownership interest in PacingCure BV. Other authors declare no competing interests.

## Acknowledgments

The authors would kindly like to acknowledge the significant contribution of Corrie Gier de Vries in the preparation of tissue sections for subsequent immunohistochemistry and the AAV Vector unit at ICGEB Trieste for producing the AAV9-hTNNT2-GFP and AAV9-hTNNT2-TBX5 viruses.

---

**Supplementary Figure 1.**
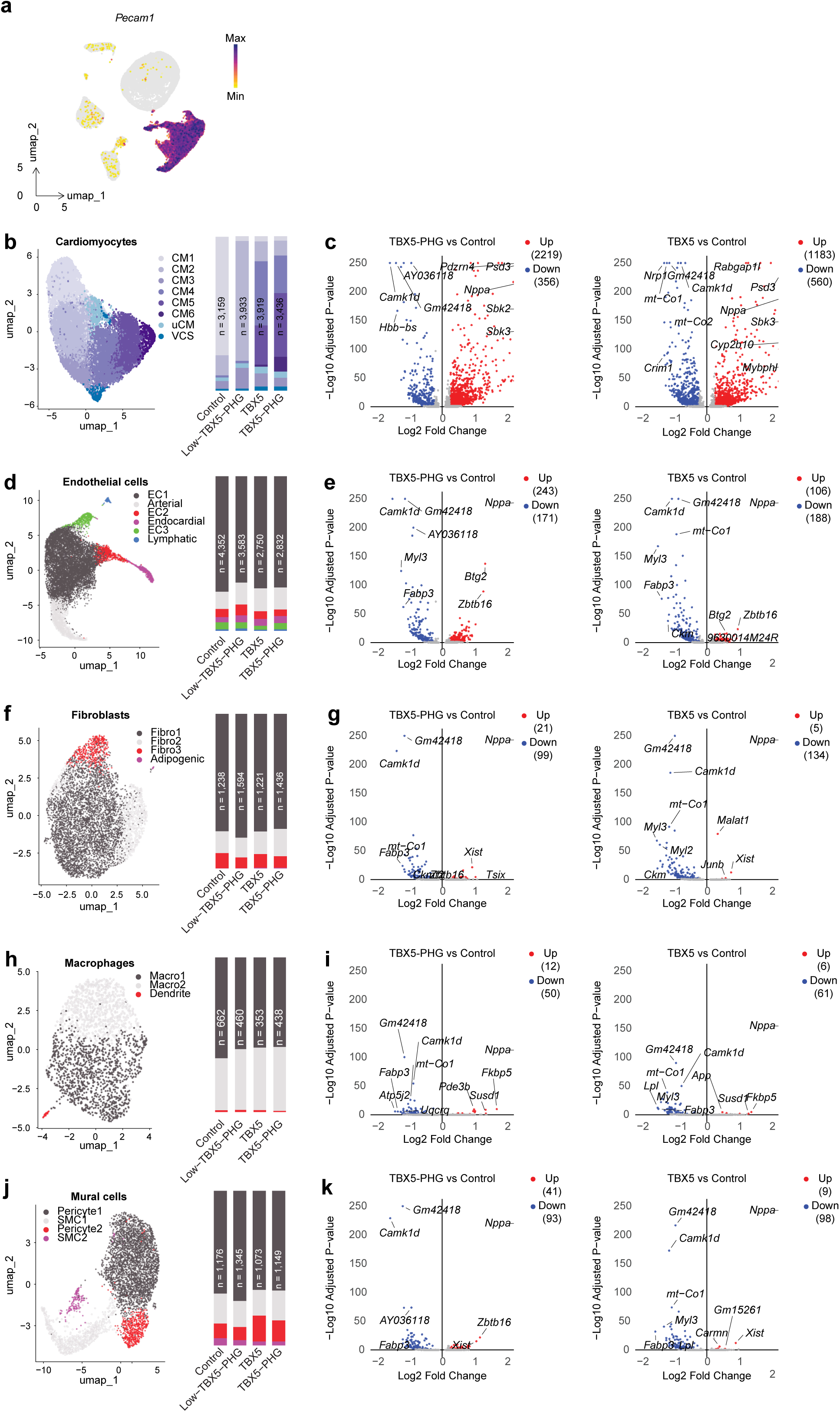
CM-specific TBX5 delivery minimally affect other cardiac cell types. (**a**) Feature plot showing *Pecam1* expression on the UMAP of all detected cell types, Composition of detected subclusters per condition (right) of (**b**) CMs (**d**) endothelial cells (**f**) fibroblasts (**h**) macrophages (**j**) mural cells, Volcano plot of DEGs (P_adj_<0.05) between AAV9-Control and AAV9-TBX5-PHG conditions (left) or between AAV9-Control and AAV9-TBX5 conditions (right) in (**c**) CM, (**e**) endothelial cells, (**g**) fibroblasts, (**i**) macrophages, (**k**) mural cells.

**Supplementary Figure 2.**
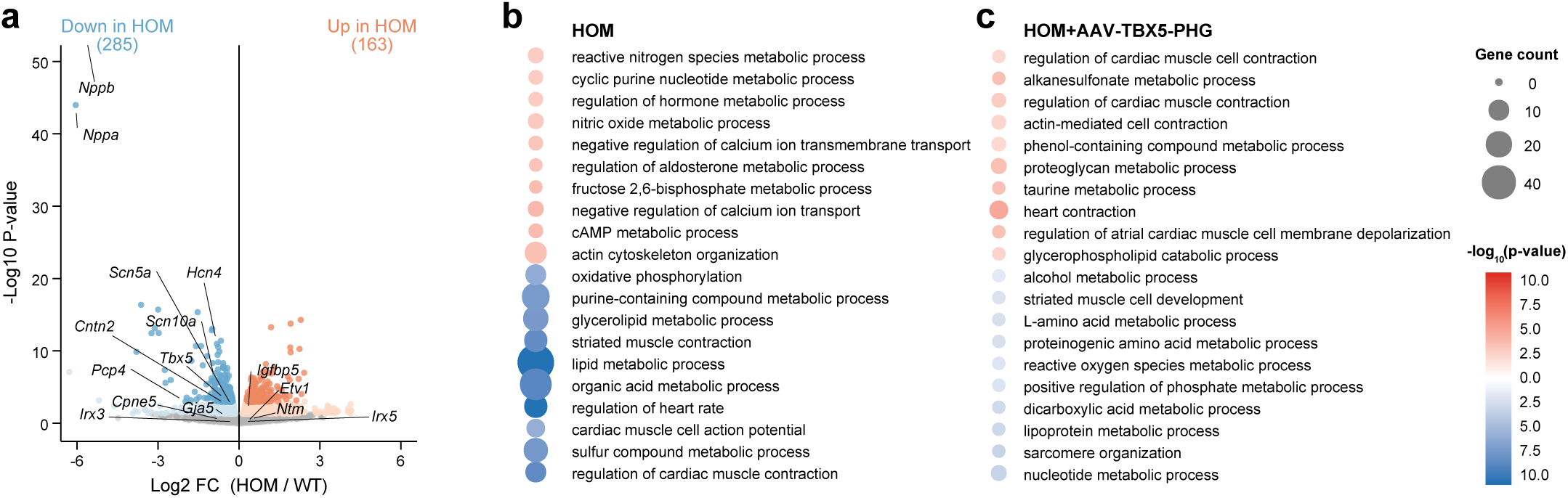
Comparison of TBX5-driven atrial and ventricular transcriptional programs. (**a**) Volcano plot of DEGs (P_adj_<0.05) between adult ventricular tissue of HOM (*Nppa-Nppb^⁻/⁻^*) and WT mice, (**b-c**) Enriched biological process Gene Ontology (GO) terms for selected gene sets. Dot plot shows GO terms enriched among the indicated gene sets. 10 GO terms per set, combining cardiac-relevant terms and additional significant terms. (**b**) terms enriched among genes upregulated (KO up) or downregulated (KO down) in *Nppa-Nppb^⁻/⁻^* hearts compared to wild-type, (**c**) terms enriched among genes upregulated or downregulated following AAV9-TBX5 administration in *Nppa-Nppb^⁻/⁻^* hearts. Dot color reflects significance (red, upregulated; blue, downregulated), and dot size indicates the number of genes from the input set annotated to each term.

**Supplementary Figure 3.**
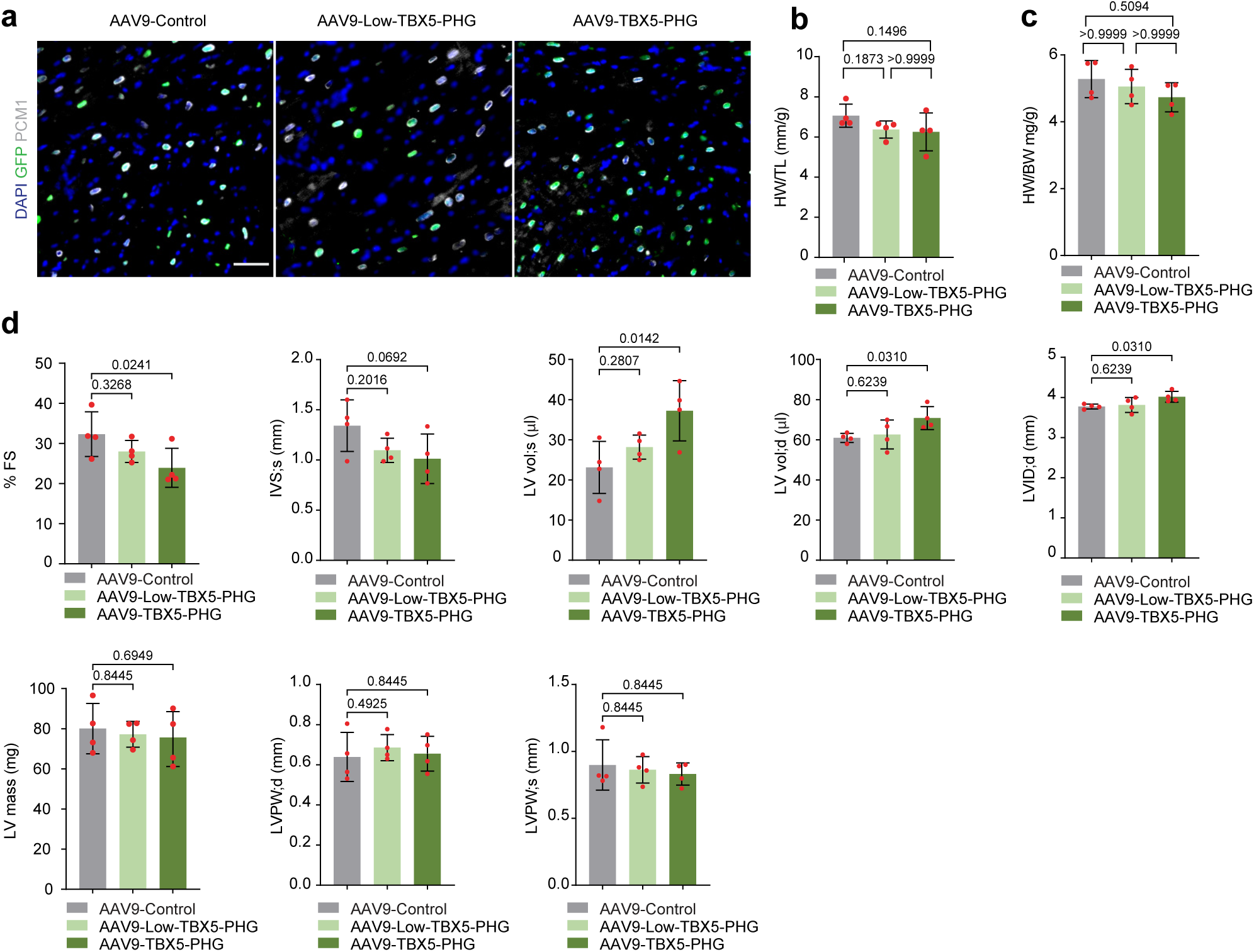
Transduction and functional profile of low and high TBX5 doses. (**a**) Immunostaining for GFP (green), PCM1 (gray) and DAPI (blue) in the ventricles of juvenile AAV9-Control, AAV9-Low-TBX5-PHG, AAV9-TBX5-PHG-injected mice (scale bar: 20 μm), (**b**) Heart weight to tibia length ratio (HW/TL) and (**c**) Heart weight to body weight ratio (HW/BW) of AAV9-Control, AAV9-Low-TBX5-PHG and AAV9-TBX5-PHG-treated mice (n=4) (**d**) Echocardiography measurements at systole (s) and diastole (d) for fraction shortening (FS), thickness (IVS;s), left ventricular internal diameter (LVID;d), LV mass, left posterior wall thickness (LVPW;s, LVPW;d) and left ventricular volume (LVvol;s, LVvol;d) in AAV9-Control, AAV9-Low-TBX5-PHG, AAV9-TBX5-PHG-injected mice (n=4). Comparison between more groups were analyzed by Kruskal-Wallis.

**Supplementary Figure 4.**
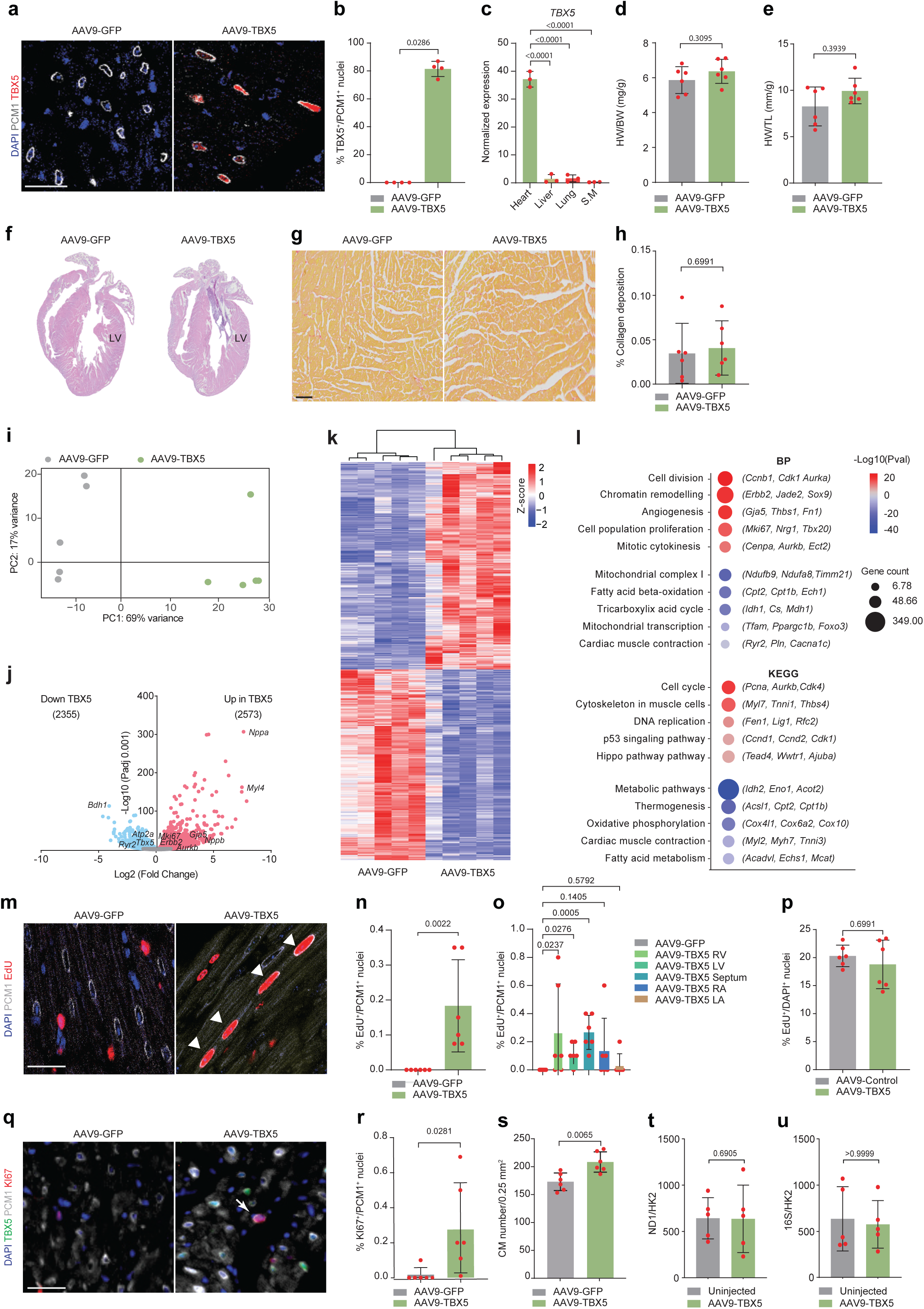
CM-specific TBX5 delivery alters the transcriptional profile of the postnatal ventricles and induces CM cell cycle re-entry. (**a**) Immunostaining for PCM1 (gray), TBX5 (red) and DAPI (blue) in the ventricles of AAV9-Control and AAV9-TBX5-treated mice (scale bar: 25 μm), (**b**) Quantification of TBX5^+^/PCM1^+^ nuclei (n=4), (**c**) TBX5 RNA expression levels in heart, liver, lung and skeletal muscle (S.K) of AAV9-TBX5 injected mice (n=3), (**d**) Heart weight to body weight (HW/BW) and (**e**) Heart weight to tibia length (HW/TL) ratio of AAV9-Control and AAV9-TBX5-treated mice (n=6), (**f**) Hematoxylin eosin staining and (**g**) picrosirius red staining of AAV9-Control and AAV9-TBX5-treated mice (scale bar: 500 μm), (**h**) Quantification of collagen deposition in AAV9-Control and AAV9-TBX5-treated mice (n=6), (**i**) Principal component analysis (PCA plot) of bulk RNA sequencing for AAV9-Control and AAV9-TBX5-treated mice, (**j**) Volcano plot of DEGs (P_adj_<0.001) in AAV9-Control and AAV9-TBX5-treated mice 14 days after injection, (**k**) Heatmap showing the normalized expression of DEGs in AAV9-Control and AAV9-TBX5 groups, (**l**) Gene ontology analysis showing biological processes (BP) and KEGG pathways based on DEGs (**Supplementary Figure 4j**), (**m**) Representative immunofluorescence staining of PCM1 (gray), EdU (red), and DAPI (blue) in ventricles from AAV9-Control- and AAV9-TBX5-treated mice. (Scale bar: 25 μm), (**n-o**) Quantification of EdU^+^/PCM1^+^ nuclei per (**n**) heart or (**o**) chamber (n=6). No EdU^+^/PCM1^+^ nuclei were detected in AAV9-control; all chambers were therefore pooled in the quantification graph, (**p**) Quantification of EdU^+^/DAPI^+^ nuclei in the ventricles of AAV9-Control and AAV9-TBX5-treated mice (n=6), (**q**) Representative immunofluorescence staining of PCM1 (gray), TBX5 (green), KI67 (red), and DAPI (blue) in ventricles from AAV9-GFP (Control)- and AAV9-TBX5-treated mice, (**r**) Quantification of KI67^+^/PCM1^+^ nuclei in the ventricles of AAV9-Control and AAV9-TBX5-treated mice (n=6), (**s**) Quantification of CM number per 0.25 mm^2^ in the ventricles of AAV9-Control and AAV9-TBX5-treated mice (n=6), (**t-u**) Normalized expression of the mitochondrial genes ND1 (**t**) and 16S (**u**) was calculated relative to the nuclear gene hexokinase (HK2), quantifying mtDNA copy number in uninjected and AAV9-TBX5-treated mice (n=5). Comparisons between two groups were analyzed using the Mann-Whitney U test. Comparisons between three and more groups were analyzed using One-Way ANOVA.

## Bibliography

1. Kim, S. & Wysocka, J. Deciphering the multi-scale, quantitative cis-regulatory code. Mol. Cell 83, 373–392 (2023).

2. Spitz, F. & Furlong, E. E. M. Transcription factors: from enhancer binding to developmental control. Nat. Rev. Genet. 13, 613–626 (2012).

3. Blassberg, R. et al. Sox2 levels regulate the chromatin occupancy of WNT mediators in epiblast progenitors responsible for vertebrate body formation. Nat. Cell Biol. 24, 633–644 (2022).

4. Chen, A. I., De Nooij, J. C. & Jessell, T. M. Graded Activity of Transcription Factor Runx3 Specifies the Laminar Termination Pattern of Sensory Axons in the Developing Spinal Cord. Neuron 49, 395–408 (2006).

5. Domingo, J. et al. Nonlinear transcriptional responses to gradual modulation of transcription factor dosage. eLife 13, RP100555 (2026).

6. Jacob, J. et al. Retinoid Acid Specifies Neuronal Identity through Graded Expression of Ascl1. Curr. Biol. 23, 412–418 (2013).

7. Kathiriya, I. S. et al. A disrupted compartment boundary underlies abnormal cardiac patterning and congenital heart defects. Nat. Cardiovasc. Res. 5, 67–83 (2026).

8. Liu, W. et al. Dissecting the impact of transcription factor dose on cell reprogramming heterogeneity using scTF-seq. Nat. Genet. 57, 2522–2535 (2025).

9. Mori, A. D. et al. Tbx5-dependent rheostatic control of cardiac gene expression and morphogenesis. Dev. Biol. 297, 566–586 (2006).

10. Naqvi, S. et al. Precise modulation of transcription factor levels identifies features underlying dosage sensitivity. Nat. Genet. 55, 841–851 (2023).

11. Ni, Z., Zhou, X.-Y., Aslam, S. & Niu, D.-K. Characterization of Human Dosage-Sensitive Transcription Factor Genes. Front. Genet. 10, 1208 (2019).

12. Seidman, J. G. & Seidman, C. Transcription factor haploinsufficiency: when half a loaf is not enough. J. Clin. Invest. 109, 451–455 (2002).

13. Van Der Lee, R., Correard, S. & Wasserman, W. W. Deregulated Regulators: Disease-Causing cis Variants in Transcription Factor Genes. Trends Genet. 36, 523–539 (2020).

14. Abramov, S. et al. Landscape of allele-specific transcription factor binding in the human genome. Nat. Commun. 12, 2751 (2021).

15. Maurano, M. T. et al. Systematic Localization of Common Disease-Associated Variation in Regulatory DNA. Science 337, 1190–1195 (2012).

16. The 1000 Genomes Project Consortium et al. A global reference for human genetic variation. Nature 526, 68–74 (2015).

17. Yan, J. et al. Systematic analysis of binding of transcription factors to noncoding variants. Nature 591, 147–151 (2021).

18. Gupta, R., Karczewski, K. J., Howrigan, D., Neale, B. M. & Mootha, V. K. Human genetic analyses of organelles highlight the nucleus in age-related trait heritability. eLife 10, e68610 (2021).

19. Miyazawa, K. et al. Cross-ancestry genome-wide analysis of atrial fibrillation unveils disease biology and enables cardioembolic risk prediction. Nat. Genet. 55, 187–197 (2023).

20. Mostafavi, H., Spence, J. P., Naqvi, S. & Pritchard, J. K. Systematic differences in discovery of genetic effects on gene expression and complex traits. Nat. Genet. 55, 1866–1875 (2023).

21. Roselli, C. et al. Multi-ethnic genome-wide association study for atrial fibrillation. Nat. Genet. 50, 1225–1233 (2018).

22. The BIOS Consortium et al. Disease variants alter transcription factor levels and methylation of their binding sites. Nat. Genet. 49, 131–138 (2017).

23. Pulice, J. L. & Meyerson, M. Amplified dosage of the NKX2-1 lineage transcription factor controls its oncogenic role in lung adenocarcinoma. Mol. Cell 85, 1311–1329.e16 (2025).

24. Steimle, J. D. & Moskowitz, I. P. TBX5: A Key Regulator of Heart Development. Curr. Top. Dev. Biol. 122, 195–221 (2017).

25. Basson, C. T. et al. Mutations in human cause limb and cardiac malformation in Holt-Oram syndrome. Nat. Genet. 15, 30–35 (1997).

26. Bruneau, B. G. et al. A Murine Model of Holt-Oram Syndrome Defines Roles of the T-Box Transcription Factor Tbx5 in Cardiogenesis and Disease. Cell 106, 709–721 (2001).

27. Li, Q. Y. et al. Holt-Oram syndrome is caused by mutations in TBX5, a member of the Brachyury (T) gene family. Nat. Genet. 15, 21–29 (1997).

28. Vanlerberghe, C. et al. Holt-Oram syndrome: clinical and molecular description of 78 patients with TBX5 variants. Eur. J. Hum. Genet. 27, 360–368 (2019).

29. Arnolds, D. E. et al. TBX5 drives Scn5a expression to regulate cardiac conduction system function. J. Clin. Invest. 122, 2509–2518 (2012).

30. Burnicka-Turek, O. et al. Coordinated Tbx3/Tbx5 transcriptional control of the adult ventricular conduction system. eLife 13, RP102027 (2025).

31. Dai, W. et al. A calcium transport mechanism for atrial fibrillation in Tbx5-mutant mice. eLife 8, e41814 (2019).

32. Kathiriya, I. S. et al. Reduced TBX5 dosage undermines developmental control of atrial cardiomyocyte identity in a model of human atrial disease. Development 153, dev205173 (2026).

33. Nadadur, R. D. et al. Pitx2 modulates a Tbx5-dependent gene regulatory network to maintain atrial rhythm. Sci. Transl. Med. 8, 354ra115 (2016).

34. Rathjens, F. S. et al. Preclinical evidence for the therapeutic value of TBX5 normalization in arrhythmia control. Cardiovasc. Res. 117, 1908–1922 (2021).

35. Smemo, S. et al. Regulatory variation in a TBX5 enhancer leads to isolated congenital heart disease. Hum. Mol. Genet. 21, 3255–3263 (2012).

36. Sweat, M. E. et al. Tbx5 maintains atrial identity in postnatal cardiomyocytes by regulating an atrial-specific enhancer network. Nat. Cardiovasc. Res. 2, 881–898 (2023).

37. Kimura, M., Kikuchi, A., Ichinoi, N. & Kure, S. Novel TBX5 Duplication in a Japanese Family with Holt–Oram Syndrome. Pediatr. Cardiol. 36, 244–247 (2015).

38. Patel, C., Silcock, L., McMullan, D., Brueton, L. & Cox, H. TBX5 intragenic duplication: a family with an atypical Holt-Oram syndrome phenotype. Eur. J. Hum. Genet. EJHG 20, 863–869 (2012).

39. Postma, A. V. et al. A Gain-of-Function TBX5 Mutation Is Associated With Atypical Holt–Oram Syndrome and Paroxysmal Atrial Fibrillation. Circ. Res. 102, 1433–1442 (2008).

40. van Ouwerkerk, A. F. et al. Patient-Specific TBX5-G125R Variant Induces Profound Transcriptional Deregulation and Atrial Dysfunction. Circulation 145, 606–619 (2022).

41. Bosada, F. M. et al. An atrial fibrillation-associated regulatory region modulates cardiac Tbx5 levels and arrhythmia susceptibility. eLife 12, e80317 (2023).

42. Man, J. C. K. et al. Genetic Dissection of a Super Enhancer Controlling the *Nppa-Nppb* Cluster in the Heart. Circ. Res. 128, 115–129 (2021).

43. Schindelin, J. et al. Fiji: an open-source platform for biological-image analysis. Nat. Methods 9, 676–682 (2012).

44. Schmidt, U., Weigert, M., Broaddus, C. & Myers, G. Cell Detection with Star-Convex Polygons. in Medical Image Computing and Computer Assisted Intervention – MICCAI 2018 (eds Frangi, A. F., Schnabel, J. A., Davatzikos, C., Alberola-López, C. & Fichtinger, G.) vol. 11071 265–273 (Springer International Publishing, Cham, 2018).

45. Afgan, E. et al. The Galaxy platform for accessible, reproducible and collaborative biomedical analyses: 2018 update. Nucleic Acids Res. 46, W537–W544 (2018).

46. Love, M. I., Huber, W. & Anders, S. Moderated estimation of fold change and dispersion for RNA-seq data with DESeq2. Genome Biol. 15, 550 (2014).

47. Sherman, B. T., Panzade, G., Imamichi, T. & Chang, W. DAVID Ortholog: an integrative tool to enhance functional analysis through orthologs. Bioinformatics 40, btae615 (2024).

48. Zheng, G. X. Y. et al. Massively parallel digital transcriptional profiling of single cells. Nat. Commun. 8, 14049 (2017).

49. Satija, R., Farrell, J. A., Gennert, D., Schier, A. F. & Regev, A. Spatial reconstruction of single-cell gene expression data. Nat. Biotechnol. 33, 495–502 (2015).

50. McGinnis, C. S., Murrow, L. M. & Gartner, Z. J. DoubletFinder: Doublet Detection in Single-Cell RNA Sequencing Data Using Artificial Nearest Neighbors. Cell Syst. 8, 329–337.e4 (2019).

51. Choudhary, S. & Satija, R. Comparison and evaluation of statistical error models for scRNA-seq. Genome Biol. 23, 27 (2022).

52. Hafemeister, C. & Satija, R. Normalization and variance stabilization of single-cell RNA-seq data using regularized negative binomial regression. Genome Biol. 20, 296 (2019).

53. Phipson, B. et al. propeller: testing for differences in cell type proportions in single cell data. Bioinformatics 38, 4720–4726 (2022).

54. Hao, Y. et al. Integrated analysis of multimodal single-cell data. Cell 184, 3573–3587.e29 (2021).

55. Dolgalev, I. Babelgene: Gene Orthologs for Model Organisms in a Tidy Data Format. (2022).

56. Gu, Z., Eils, R. & Schlesner, M. Complex heatmaps reveal patterns and correlations in multidimensional genomic data. Bioinformatics 32, 2847–2849 (2016).

57. Street, K. et al. Slingshot: cell lineage and pseudotime inference for single-cell transcriptomics. BMC Genomics 19, 477 (2018).

58. Van den Berge, K., et al. Trajectory-based differential expression analysis for single-cell sequencing data. Nat. Commun. 11, 1201 (2020).

59. Yu, G., Wang, L.-G., Han, Y. & He, Q.-Y. clusterProfiler: an R Package for Comparing Biological Themes Among Gene Clusters. OMICS J. Integr. Biol. 16, 284–287 (2012).

60. Quiros, P. M., Goyal, A., Jha, P. & Auwerx, J. Analysis of mtDNA/nDNA Ratio in Mice. Curr. Protoc. Mouse Biol. 7, 47–54 (2017).

61. Giovou, A. E., et al. Nppa and Nppb Deficiency Drives Ventricular Hypertrophy and Subendocardial Gene Deregulation in the Mouse Heart. Int. J. Mol. Sci. 27, 2450 (2026).

62. Cao, Y. et al. In Vivo Dissection of Chamber-Selective Enhancers Reveals Estrogen-Related Receptor as a Regulator of Ventricular Cardiomyocyte Identity. Circulation 147, 881–896 (2023).

63. Mulleners, O. J., Van Der Maarel, L. E., Christoffels, V. M. & Jensen, B. The trabecular and compact myocardium of adult vertebrate ventricles are transcriptionally similar despite morphological differences. Ann. N. Y. Acad. Sci. 1545, 76–90 (2025).

64. Lonsdale, J. et al. The Genotype-Tissue Expression (GTEx) project. Nat. Genet. 45, 580–585 (2013).

65. Oh, Y. et al. Transcriptional regulation of the postnatal cardiac conduction system heterogeneity. Nat. Commun. 15, 6550 (2024).

66. Ferreira, J. P., Overton, K. W. & Wang, C. L. Tuning gene expression with synthetic upstream open reading frames. Proc. Natl. Acad. Sci. U. S. A. 110, 11284–11289 (2013).

67. Wang, X.-Q. & Rothnagel, J. A. 5′-Untranslated regions with multiple upstream AUG codons can support low-level translation via leaky scanning and reinitiation. Nucleic Acids Res. 32, 1382–1391 (2004).

68. Goodyer, W. R. et al. Transcriptomic Profiling of the Developing Cardiac Conduction System at Single-Cell Resolution. Circ. Res. 125, 379–397 (2019).

69. Ahmadian, M. et al. PPARγ signaling and metabolism: the good, the bad and the future. Nat. Med. 19, 557–566 (2013).

70. Chandra, M., Miriyala, S. & Panchatcharam, M. PPARγ and Its Role in Cardiovascular Diseases. PPAR Res. 2017, 6404638 (2017).

71. Vega, R. B. & Kelly, D. P. Cardiac nuclear receptors: architects of mitochondrial structure and function. J. Clin. Invest. 127, 1155–1164 (2017).

72. Friedrich, F. W. et al. FHL2 expression and variants in hypertrophic cardiomyopathy. Basic Res. Cardiol. 109, 451 (2014).

73. Prosdocimo, D. A. et al. KLF15 and PPARα Cooperate to Regulate Cardiomyocyte Lipid Gene Expression and Oxidation. PPAR Res. 2015, 201625 (2015).

74. Hong, J.-H. & Zhang, H.-G. Transcription Factors Involved in the Development and Prognosis of Cardiac Remodeling. Front. Pharmacol. 13, (2022).

75. Churko, J. M. et al. Defining human cardiac transcription factor hierarchies using integrated single-cell heterogeneity analysis. Nat. Commun. 9, 4906 (2018).

76. Cao, L. et al. PGC-1α: key regulator of mitochondrial biogenesis and cellular differentiation in metabolic and regenerative tissues. Cell Biosci. 16, 9 (2025).

77. Miquerol, L. et al. Resolving cell lineage contributions to the ventricular conduction system with a Cx40-GFP allele: A dual contribution of the first and second heart fields. Dev. Dyn. 242, 665–677 (2013).

78. Pallante, B. A. et al. Contactin-2 Expression in the Cardiac Purkinje Fiber Network. Circ. Arrhythm. Electrophysiol. 3, 186–194 (2010).

79. Nguyen, P. D. et al. Interplay between calcium and sarcomeres directs cardiomyocyte maturation during regeneration. Science 380, 758–764 (2023).

80. Kathiriya, I. S. et al. Modeling Human TBX5 Haploinsufficiency Predicts Regulatory Networks for Congenital Heart Disease. Dev. Cell 56, 292–309.e9 (2021).

81. Steimle, J. D. et al. Single-nuclei transcriptomics reveals TBX5-dependent targets in a patient with Holt-Oram syndrome. J. Clin. Invest. 135, e180670 (2025).

82. Sweat, M. E. et al. TBX5 and CHD4 Coordinately Activate Atrial Cardiomyocyte Genes to Maintain Cardiac Rhythm Homeostasis. Circulation 152, 784–801 (2025).

83. Pezhouman, A. et al. Transcriptional, Electrophysiological, and Metabolic Characterizations of hESC-Derived First and Second Heart Fields Demonstrate a Potential Role of TBX5 in Cardiomyocyte Maturation. Front. Cell Dev. Biol. 9, 787684 (2021).

84. Froese, N. et al. Hypoxia Attenuates Pressure Overload-Induced Heart Failure. J. Am. Heart Assoc. 13, e033553 (2024).

85. Man, J. C. K. et al. Variant Intronic Enhancer Controls SCN10A-short Expression and Heart Conduction. Circulation 144, 229–242 (2021).

86. Sotoodehnia, N. et al. Common variants in 22 loci are associated with QRS duration and cardiac ventricular conduction. Nat. Genet. 42, 1068–1076 (2010).

87. Shekhar, A. et al. Transcription factor ETV1 is essential for rapid conduction in the heart. J. Clin. Invest. 126, 4444–4459 (2016).

88. Aharonov, A. et al. ERBB2 drives YAP activation and EMT-like processes during cardiac regeneration. Nat. Cell Biol. 22, 1346–1356 (2020).

89. Bouwman, M. et al. Cross-species comparison reveals that Hmga1 reduces H3K27me3 levels to promote cardiomyocyte proliferation and cardiac regeneration. Nat. Cardiovasc. Res. 4, 64–82 (2025).

90. Li, X. et al. Inhibition of fatty acid oxidation enables heart regeneration in adult mice. Nature 622, 619–626 (2023).

91. Monroe, T. O. et al. YAP Partially Reprograms Chromatin Accessibility to Directly Induce Adult Cardiogenesis In Vivo. Dev. Cell 48, 765–779.e7 (2019).

92. Siatra, P. et al. Return of the Tbx5; lineage-tracing reveals ventricular cardiomyocyte-like precursors in the injured adult mammalian heart. Npj Regen. Med. 8, 13 (2023).

93. Shakked, A. et al. Redifferentiated cardiomyocytes retain residual dedifferentiation signatures and are protected against ischemic injury. Nat. Cardiovasc. Res. 2, 383–398 (2023).

